# RNase H1 counteracts DNA damage and ameliorates SMN-dependent phenotypes in a *Drosophila* model of Spinal Muscular Atrophy

**DOI:** 10.1101/2025.04.17.649348

**Authors:** Livia Scatolini, Lucia Graziadio, Damiano Guerrini, Paolo Maccallini, Carmen Marino, Raffaella di Vito, Amber Hassan, Manuela Grimaldi, Marco Fidaleo, Roozbeh Dehghannasiri, Gabriele Proietti, Roberto Piergentili, Erica Salvati, Stefano Cacchione, Cristiano De Pittà, Ylli Doksani, Anna Maria D’Ursi, James Wakefield, Lu Chen, Maurizio Gatti, Alessio Colantoni, Alessandro Usiello, Grazia D. Raffa

## Abstract

Spinal Muscular Atrophy (SMA) is caused by a deficiency of the Survival Motor Neuron (SMN) protein. Mutations in SMN disrupt mRNA splicing and translation, leading to maladaptive changes in transcriptomes, proteomes, neuroinflammation, and metabolism, which drive motor neuron degeneration in SMA patients. Using a *Drosophila* SMA model, we found that systemic depletion of Smn leads to accumulation of RNA:DNA hybrids (R-loops), increased DNA damage, dysregulation of amino acids and sugar metabolism and activation of the innate immune response, recapitulating key pathological features reported in mammalian models and severe SMA patients. Persistent DNA damage in Smn-deficient flies alters cell proliferation rates in larval brains and induces extensive cell death in the developing eye. Importantly here, we show that stimulating the resolution of RNA:DNA hybrids with transgenic human RNAse H1 prevents the accumulation of DNA damage and attenuates the transcriptome and amino acid alterations induced by Smn depletion, mitigating the Smn-dependent cellular and developmental abnormalities, in Smn-deficient flies. Our data suggest that depletion of Smn causes an accumulation of aberrant transcripts and chronic DNA damage, which—along with the altered metabolomic profiles associated with Smn deficiency—trigger systemic inflammatory responses, ultimately affecting neuronal function and survival.

## INTRODUCTION

Spinal Muscular Atrophy (SMA) is a devastating neuromuscular disorder, characterized by progressive motoneuron dysfunction and death^1^. SMA is caused by the loss of the Survival Motor Neuron (SMN) protein, which is part of a complex that mediates the assembly of small nuclear RNAs (snRNAs) into ribonucleoproteins (snRNPs) that are responsible for precursor mRNA splicing ^2, 3^. Nascent snRNAs, transcribed by RNA polymerase II, are modified at their 5′ ends, by the addition of an m7-monomethylguanosine cap (m7G/MMG cap); snRNAs precursors are subsequently translocated into the cytoplasm and bind to the Gemin5 subunit of the SMN complex ^4^, which mediates snRNP assembly with the Sm core proteins ^5^. SMN physically interacts with the trimethylguanosine synthase 1, TGS1 ^6, 7,4, 5^, which converts the MMG cap of snRNAs into a 2,2,7-trimethylguanosine cap (TMG cap).

Precise formation of the 3′ ends of snRNAs occurs through cleavage of the precursors by the Integrator complex ^8^, followed by trimming by the TOE1 deadenylase ^9^. The 3′ end processing and quality control of snRNAs requires cap hypermethylation of their precursors by TGS1 ^10, 11^, which also aids in pre-snRNP trafficking ^12, 13^ to the nuclear Cajal bodies, for their final maturation ^14, 15^.

The chain of events responsible for the death of SMN-deficient motoneurons is still poorly understood. Reduced SMN levels result in defective snRNP assembly, impair the splicing of precursor mRNAs essential for motoneuron survival, and cause extensive transcriptome alterations in SMA models ^2, 16, 17, 18, 19, 20^. *SMN* mutations also affect the maturation of histone mRNAs through U7 snRNP ^21, 22^, ribosome priming and protein synthesis ^23, 24^. Furthermore, SMN plays additional roles in motoneurons, including axonal transport ^25^ and neuromuscular junction organization ^22, 26^. More recently, compelling findings highlighted that activation of the innate immune response and neuroinflammation contribute to pathogenesis in both SMA patients and animal models ^27, 28, 29, 30,31^. Indeed, SMA1 patients show elevated levels of proinflammatory cytokines in their cerebrospinal fluid, which were partially decreased after treatment with Nusinersen, an antisense oligonucleotide (ASO) -based therapy that increases the production of the full-length SMN protein ^27^. However, the relative contribution of SMN-related processes to the activation of the immune response is still unclear. Additionally, consistent with the multisystemic nature of SMA^32^, the loss of SMN in both animal models and SMA patients elicits severe deregulation of metabolic pathways related to cellular energy homeostasis ^33, 34, 35, 36^ and metabolism of nucleotides, lipids and amino acids ^37, 38, 39, 40^.

SMN also preserves genome integrity, by preventing the accumulation of RNA:DNA hybrids (R-loops) in transcription termination regions and ribosomal DNA ^41, 42, 43, 44, 45, 46, 47^, hence alterations in the splicing process can increase R-loops formation, causing DNA damage and genome instability ^48, 49, 50, 51, 52, 53, 54^. In *Drosophila,* the *SMN* and *TGS1* orthologs (*Smn* and *Tgs1*) cooperate in snRNP biogenesis and *Tgs1* overexpression ameliorates the *Smn* loss-of-function phenotypes and vice versa ^7, 55^. Accordingly, loss of either SMN or TGS1 in HeLa cells induces the formation of mRNAs that carry alterations in splicing and 3′ readthrough extensions ^10^, thus highlighting a role for the SMN-TGS1 axis in regulating molecular processes crucial for splicing fidelity, neuron stability, and relevant for modeling SMA pathogenesis.

Using a *Drosophila* SMA model based on ubiquitous depletion of Smn ^56^, we investigated the involvement of R-loop metabolism in the cellular and inflammatory phenotypes caused by loss of Smn. Our findings indicate that Smn depletion results in snRNP instability and transcriptional stress, which can lead to the accumulation of RNA:DNA hybrids, chronic DNA damage, inflammation and ultimately to a decline in neuron function and survival.

## RESULTS

### Innate immune response and metabolic alterations in *Smn*-deficient flies

To obtain severe depletion of *Smn*, we generated *ActGAL4>SmnRNAi; Smn^X^*^7^/+ flies, (henceforth designated as *SmnRNAi*), which are heterozygous for a deletion of the *Smn* locus (*Smn^x7^*) and bear the *Actin-GAL4* driver in combination with an inducible *UAS-SmnRNAi* construct. *SmnRNAi* flies die as early pupae and have been previously used to model SMA-relevant phenotypes ^56^. RT-qPCR performed on RNA obtained from either *SmnRNAi* or control larvae bearing the driver alone *(Actin-GAL4/+;* henceforth designated as CTR) as well as on RNA from brains of the same larvae, showed that the abundance of the *Smn* mRNA was reduced by 79% and 88%, respectively, compared to controls (Figure 1A).

**Figure 1.**
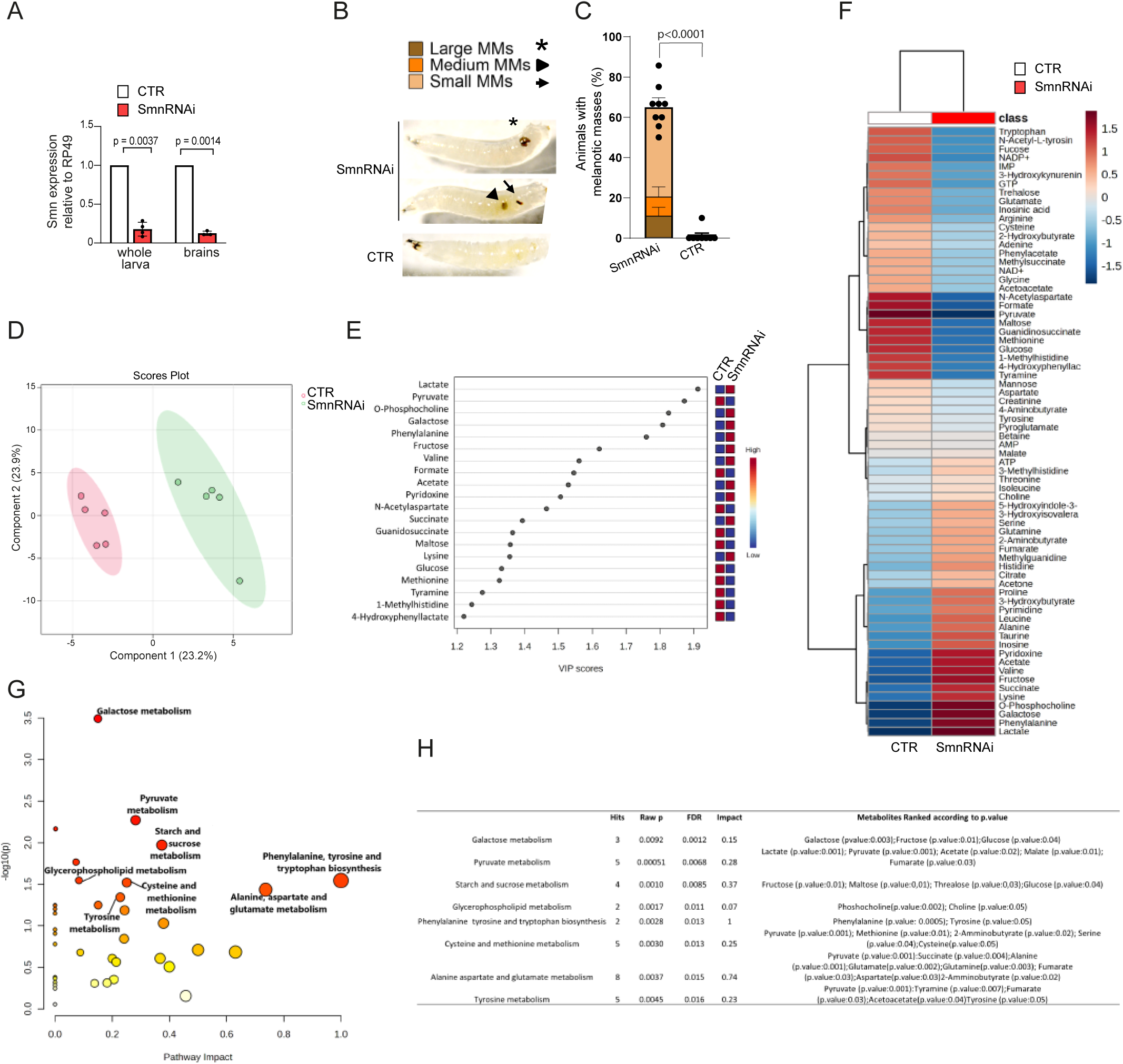
**Metabolic alterations and melanization response in *SmnRNAi* flies** (A) Efficiency of RNAi-mediated *Smn*-knockdown. RT-qPCR measurements of *Smn* transcripts levels in third instar larvae or brains, with the indicated genotypes. Bars represent the fold changes of *Smn* transcript levels normalized to *Rp49* in *SmnRNAi vs* control samples (CTR; set to 1). Bars: mean, SD; n in bars = number of independent experimental replicates (larvae n=4; brains n=3; average of 3 technical replicates per samples p values: two-tailed unpaired t-test, df=3 (larvae); df=2 (brains). (B) Representative images of melanotic masses of different size in *SmnRNAi* larvae; the frequencies of these masses are quantified in (C). (C) Melanotic masses (MM) in third instar *SmnRNAi* larvae increase in number and size over time; >200 wandering larvae for each genotype were scored by visual inspection up to 24 hours after collection in 9 experiments. Bars: mean of the average number of MM (of any size) per sample + SEM; p values: two-tailed unpaired t-test, df=8. (D) PLS-DA score scatter plots showing the metabolomic profiles of *SmnRNAi* vs CTR larvae. The cluster analyses are reported in the Cartesian space that is described by the principal components PC1 and PC2. Partial Least Squares Discriminant Analysis (PLS-DA) was evaluated using cross-validation (CV) analysis; CV tests, 1,0 accuracy values on PC1 and PC2, respectively, and positive 0.60 and 0.75 Q2 indexes. (E) VIP score graphs of the metabolites discriminating CTR and *SmnRNAi* metabolomic profile. Metabolites characterized by a VIP score > 1.2 are shown. (F) Hierarchical heatmap generated by MetaboAnalyst software based on the Euclidean distance and Ward’s algorithm. The heatmap is calculated based on metabolite concentrations. The bar color represents each metabolite’s abundance on a normalized scale from blue (low level) to red (high level). The dendrogram on the top is based on the similarity of the metabolomic profile relative to each sample cluster. The dendrogram on the left is based on the metabolite abundance profiles. (G) Pathway analysis shows all matched pathways according to p-values from pathway enrichment analysis (y-axis) and pathway impact values from pathway topology analysis (x-axis). The color and size of each circle are based on p-values and pathway impact values, respectively. Small p-values and large pathway impact red circles indicate that the pathway is greatly perturbed. (H) Biochemical pathways performed using MetPa. The table reports discriminating pathways of the metabolomic profiles of CTR and *SmnRNAi* larvae. The Hits are the matched number of metabolites from the user-uploaded data. P-value is calculated from the enriched analysis. False discovery rate (FDR) is the portion of false positives above the user-specified score threshold. Pathway impact is a combination of centrality and pathway enrichment results. It is calculated by adding up the importance measures of each of the matched metabolites and then dividing by the sum of the importance measures of all metabolites in each pathway. Pathways with an impact value closer to 1 are those that most discriminate the analyzed clusters.

Previous work has shown that larvae expressing Smn variants bearing substitutions in clinically relevant conserved residues, display activation of the innate immune response ^30^, accompanied by the formation of melanotic masses, tumoral formations in the larval hemolymph typically generated in response to pathogens, necrosis, damage, or genetic perturbations. We found that third instar *SmnRNAi* mutant larvae display similar melanotic masses, which varied in number and size (average: 1.1 masses per larva; Figure 1B and 1C).

Through untargeted ^1^H-NMR metabolomic analysis, we then explored whether *SmnRNAi* larvae displayed biochemical abnormalities that overlap to some extent with those recently identified in the cerebrospinal fluid (CSF) of SMA patients and in the central nervous system and liver of SMNΔ7 mice models ^37, 38, 39, 40^. The metabolomic profile of *SmRNAi* larvae identified 71 metabolites in 1D-NOESY spectra (Figure S1). Notably, partial least-squares discriminant analysis (PLS-DA) highlighted significant alterations in the metabolomic profile of *SmnRNAi* polar extracts compared to controls (Figure 1D). Specifically, Variable Importance in Projection (VIP) Analysis (Figure 1E), showed that galactose, fructose, maltose and glucose were among the analytes that differentiated *SmnRNAi* larvae from CTR larvae (Figure 1F), further supporting the involvement of SMN in regulating sugar metabolism. Consistent with this, pathway enrichment analysis revealed a prominent dysregulation of the Galactose, Starch and Sucrose metabolism in *SmnRNAi* larvae (Figure 1G).

The role of Smn levels in influencing cell bioenergetics, as reported in mammalian models and SMA patients ^57^, was confirmed in *Drosophila SmnRNAi* larvae. Indeed, PLS-DA highlighted that lactate, pyruvate, acetate and succinate were among the analytes that mostly discriminated the *SmnRNAi* and CTR metabolomic profiles (Figure 1E, F, H). Accordingly, pathway enrichment analysis revealed a remarkable alteration of pyruvate metabolism in *SmnRNAi* larvae (Figure 1G).

Metabolomic investigations also confirmed that Smn deficiency disrupts aminoacidic metabolism homeostasis, as shown by the discriminatory power of phenylalanine, valine, lysine, N-acetyl aspartate, methionine, and tyramine between *SmnRNAi* and CTR polar extracts (Figure 1F). In line with this, pathway enrichment analysis identifies phenylalanine, tyrosine and tryptophan biosynthesis, cysteine and methionine metabolism, alanine, aspartate, and glutamate metabolism among the most dysregulated pathways in *SmnRNAi* larvae (Figure 1G and 1E).

Overall, the present ^1^H-NMR metabolomic data confirms in the *Drosophila* model the impact of SMN level on regulating cellular metabolism and energy homeostasis processes previously reported in SMA patients and mammalian models ^34, 35, 36, 38, 39, 58^.

### Consequences of R-loops accumulation in *Smn*-deficient flies

Genomic DNA extracted from the larvae described above was subjected to dot blot probed with the S9.6 antibody, specific for RNA:DNA hybrid structures ^59, 60^. R-loop abundance was two-fold higher in *SmnRNAi* larvae compared to control larvae (Figures 2A and 2B).

**Figure 2.**
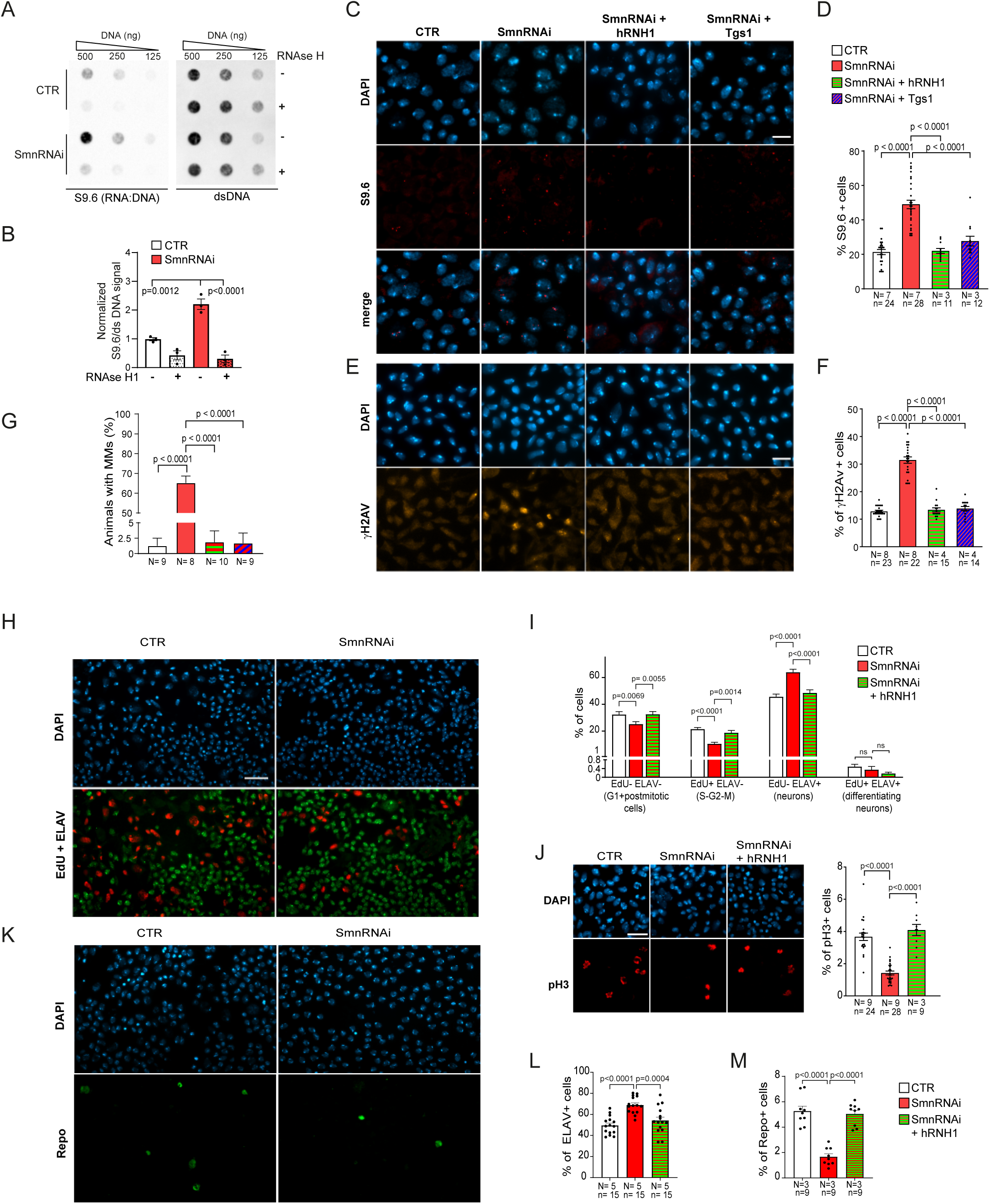
**RNA:DNA hybrids, DNA damage foci and alterations of the cell cycle profiles in *SmnRNAi* larval brains, are suppressed by overexpression of either human RNaseH1 (hRNH1) or Tgs1** (A-B) Quantification of the RNA:DNA hybrids in in wild type control (CTR) and *SmnRNAi* larvae. (A) Dot blots performed on genomic DNA extracted from whole larvae with the indicated genotypes, treated with RNase III (to reduce background signals for double-stranded RNA) according to ^112^, incubated or not with hRNH1 and probed with antibodies against RNA:DNA hybrids (S9.6, left panel) or dsDNA (as loading control, right panel). (B) Densitometric analysis of the abundance of RNA:DNA hybrid structures. Values are from 3 biological replicates and are normalized relative to the loading control. Bars: mean, SEM; p value: one-way ANOVA with Dunnett’s multiple comparisons test, F (3, 8) = 38.83. (C) Brains from larvae with the indicated genotypes (see text and Table S2A), were immunostained with the S9.6 antibody, according to ^59^, to detect RNA:DNA hybrids in nuclei, which were counterstained with DAPI. Compared to controls, *SmnRNAi* nuclei display an enrichment in S9.6 foci, quantified in (D), which is suppressed by either hRNH1 or Tgs1. Size bar in (C,E): 10µM (E) Immunostaining with the γH2Av antibody shows increased frequency of DNA damage foci in *SmnRNAi* nuclei, compared to control nuclei and to nuclei expressing either hRNH1 or Tgs1 (quantified in F). (D,F) Quantification of the S9.6 and γH2Av foci, in the IF experiments reported in (C, E). Foci were automatically quantified with the Zeiss Zen software. Bars: mean, SEM; n in bars = number of brains examined in N experiments. p values: one-way ANOVA, F (3, 71) = 40.33 (D); F (3, 70) = 133.4 (F);cells examined per sample: > 3,000 (D) and > 8,000 (F). (G) Quantification of melanotic masses; they are virtually absent in CTR, very frequent in *SmnRNAi* larvae and back to control level in *SmnRNAi+Tgs1* and *SmnRNAi + hRNH1* larvae; p values: one-way ANOVA, F (3,32) = 195.6; see also Figures 1B and 1C. (H) Brains with the indicated genotypes incubated with EdU for 2h, immunostained for ELAV and counterstained with DAPI. Size bar for all panels: 20 µM (I) Quantification of cell cycle phases distribution in CTR and *SmnRNAi* brains (Number of experiments=3; number of brains=9; > 15,000 cells examined per sample). Values are mean plus SEM; p values: two-way ANOVA with Tukey’s Multiple Comparisons test; interaction effect: F (6, 96) = 18.54, p<0.0001 (See also Figure S2). (J) Left: Brains with the indicated genotypes immunostained with anti-Phospho-Histone H3 and counterstained with DAPI. Right: Quantification of PH3-labeled mitotic cells; > 13,000 cells examined per sample. Bars: mean, SEM. N: number of experiments; n: number of brains; p values: one-way ANOVA, F (2, 56) = 53.74. (K) Brains with the indicated genotypes immunostained for Repo and counterstained with DAPI. (L) Quantification of ELAV positive cells in brains with the indicated genotypes (> 17,000 cells examined per sample). p values: one-way ANOVA, F (2, 42) = 15.39. (M) Quantification of Repo positive cells in brains with the indicated genotypes (> 14000 cells examined per sample). p values: one-way ANOVA, F (2, 24) = 41.25.

To assess whether the increased R-loop formation caused by *Smn* depletion results in the accumulation of DNA damage foci, CTR and RNAi larval brains were immunostained with either the S9.6 antibody or an antibody against the DNA damage marker γH2Av ^61^ and analyzed for the presence of S9.6 and nuclear DNA damage foci. In S*mnRNAi* brains, the S9.6 foci increased by 2.8-fold and the γH2Av foci by 2.2-fold compared to controls (Figures 2C-2F). In addition, TUNEL assay, combined with immunofluorescence staining for the pan-neuronal marker ELAV, revealed extensive DNA fragmentation signals in *SmnRNAi* brains compared to controls (Figures S2A and S2B).

To assess whether the *Smn RNAi*-induced DNA damage affects the cell cycle progression of proliferating neuroblasts, we incubated larval brains with 5-Ethynyl-2′-deoxyuridine (EdU) for 2 hours, followed by immunostaining for ELAV, to label cells in S phase and postmitotic neurons, respectively. Brains were also stained for the phospho-histone 3 (pH3) to detect mitotic cells (Figures S2C-S2G). A 2-hours EdU pulse resulted in a nearly complete labeling of the population of cells that entered the S phase during the incubation time and proceeded through mitosis, with a minimal fraction of labeled differentiating postmitotic cells (Figure S2D and S2E). Compared to controls, *SmnRNAi* brains showed a decreased proportion of S phase (EdU+ ELAV-) cycling cells, (21.4% vs 10.5%), whereas terminally differentiated cells (EdU-ELAV+) increased from 46% to 64%. *SmnRNAi* brains also showed a small but significant decrease (32% vs 25%) in the frequency of EdU-ELAV-cells, which comprise both non-dividing cells, excluding neurons, and cells in the G1 phase (Figures 2H and 2I). Accordingly, mitotic cells (pH3+) decreased from 3.7% to 1.4% (Figure 2J). Overall, in *SmnRNAi* brains, the fraction of ELAV+ cells increased to 69% compared to 49% of controls, whereas the fraction of cells labeled by the glial marker Repo was reduced from 5.3% to 1.7% (Figures 2K, 2L and 2M). These results may be indicative of loss of the proliferative potential of neuroblasts and glia cells and early senescence of the precursors, which would cause their premature differentiation. However, they may also be indicative of potential delays in the duration of S and/or G2 phases.

To assess whether DNA damage foci in *SmnRNAi* brains were a consequence of augmented R-loops, we generated *SmnRNAi + hRNH1* flies, bearing the *Actin-GAL4* driver, the UAS-*SmnRNAi* construct and a UAS-driven construct encoding human RNH1, which degrades the RNA strand in RNA:DNA hybrids ^62^. As expected, compared to *SmnRNAi* brains with no hRNH1 expression, *SmnRNAi + hRNH1* brains had a reduced number of S9.6 positive foci, comparable to that observed in control brains (Figures 2C and 2D). hRNH1 expression also reduced the DNA damage foci in *SmnRNAi* brains to control levels (Figures 2E-2F) and normalized the proportions of cells in S and M phases, as well as those of terminally differentiated ELAV+ cells and Repo+ glial cells (Figures 2I-2L). The enrichment in S9.6 and γH2Av foci in *Smn-*depleted larval brains was restored to control levels also by constitutive expression (from the tubulin promoter) of GFP-tagged hRNH1 (*SmnRNAi + hRNH1-GFP*), while the number of foci in *SmnRNAi + hRNH1^D210N^-GFP* brains, expressing catalytically inactive hRNH1, remained comparable to that observed in *SmnRNAi* brains (Figure S3A and data not shown).

We also analyzed *SmnRNAi* brains with increased Tgs1 expression, (*SmnRNAi + Tgs1*; expressing an inducible UAS-*GFP-Tgs1* construct ^7^). These brains showed a ∼40% decrease of S9.6 foci and a ∼60% decrease of γH2Av foci compared to those of *SmnRNAi* larvae (Figures 2C-2F) and a partial rescue of the proportions of ELAV+ cells and Repo+ glial cells (Figures S3B and S3C). The observation that Tgs1 alleviates the DNA damage phenotypes caused by *Smn* depletion confirms the close functional interactions between the two proteins ^7^.

*hRNH1* or *Tgs1* overexpression also rescued the melanotic tumors (or melanotic masses) phenotype caused by Smn depletion (Figure 2G). This result indicates that factors that counteract R-loops formation can also rescue the melanotic tumor phenotype likely determined by an inflammatory response to damaged DNA ^30, 63, 64^.

### Human RNH1 expression improves the eye development defects caused by Smn depletion in the eye imaginal disc

The development of the eye-antenna imaginal disc is an excellent system to investigate the consequences of neural cell death. We have previously shown that *Smn* depletion induces severe eye developmental defects that are rescued by overexpression of Tgs1^7^. To assess whether R-loop-dependent DNA damage contributes to cell death in the eye structures, we analyzed the development of the retinal neuroepithelium upon targeted knockdown of *Smn* in the larval eye imaginal discs. We generated *ey>SmnRNAi* flies expressing *UAS*-*SmnRNAi* in the eye imaginal discs (driven by *ey-Gal4*), alone or in combination with hRNH1, and evaluated eye development in both larvae and adult flies. The eye-antennal imaginal discs from *ey>SmnRNAi*, *ey>SmnRNAi + hRNH1* and control larvae were immunostained for Caspase-3 (an apoptotic marker) and ELAV, which labels the neuronal precursors of photoreceptors ^65^. The eye discs of *ey>SmnRNAi* flies displayed strong Caspase signals and reduced extension of the ELAV-stained area compared to control discs. Remarkably, both defects were partially but significantly corrected by hRNH1 expression (Figures 3A-3C).

**Figure 3.**
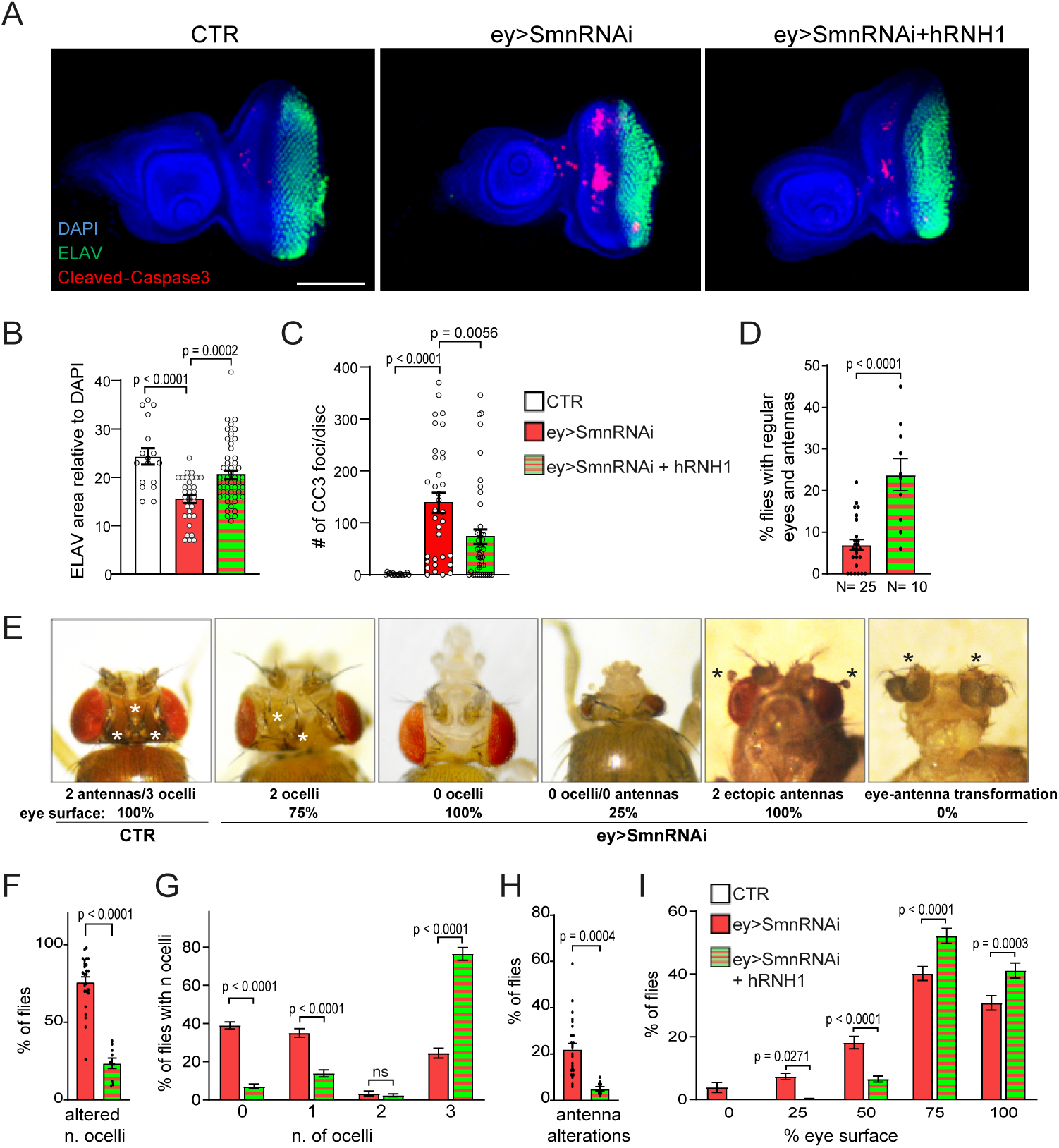
***Smn* downregulation in the eye imaginal discs induces eye and head defects that are rescued by hRNH1 expression** (A) Representative examples of eye-antennal imaginal discs from flies carrying *ey-Gal4* (*ey*) alone (CTR), ey and the *SmnRNAi* construct, or ey, *SmnRNAi* and the *hRNH1* expressing transgene. Discs were stained for ELAV (which labels developing photoreceptor cells) and Cleaved-Caspase-3 (CC3, which marks cells undergoing apoptosis) and counterstained with DAPI. Scale bar: 100 μm. (B,C) Histogram showing the quantification of ELAV (B) and CC3 (C) positive foci in imaginal discs shown in (A). Number of discs analyzed: CTR, 17; ey>*SmnRNAi*, 34; ey>*SmnRNAi+hRNH1*: 45. p values: one-way ANOVA, F (2, 93) = 14.22 (B); F (2, 93) = 12.72 (C). (D-E) *Smn* downregulation in the eye imaginal discs impairs eye development. The Histogram (D) shows the percentage of ey>*SmnRNAi* and ey>*SmnRNAi+hRNH1* flies devoid of eye and/or with head defects, with representative examples shown in (E) where white and black asterisks indicate ocelli and antennae, respectively. Bars: mean, SEM; N= number of independent experiments; the number of flies analyzed (n) is reported in (F-I); p values: two-tailed unpaired t-test, df=33 (F-I) Histogram bars representing the percentages of flies with abnormal ocelli, antennae, or defective eye surface in *ey>SmnRNAi* and *ey>SmnRNAi + hRNH1* samples. The defects in the eye and ocelli development in control flies are negligible (n>4000). *ey>SmnRNAi* flies: n= 735 for F-G; n=293 for H-I. *ey>SmnRNAi + hRNH1* flies: n= 817 for (F-G); n=507 for (H-I). p values: two-tailed unpaired t-test, df=33 in (D, F) and (H); two-way ANOVA with Šídák’s multiple comparisons test in (G) (N=13 and 11; interaction effect, F (3, 84) = 159.7; p<0.0001) and I (N=20 and 17; interaction effect, F (4, 175) = 17.64; p<0.0001).

*ey>SmnRNAi* adult flies exhibit partially reduced eyes^7^ with over 90% showing defects in eye development and alterations in the number and/or morphology of structures derived from the eye-antenna imaginal discs (Figures 3D and 3E), comprising abnormal and/or ectopic antennae, and reduced number of ocelli (ocelli derive from a distinct type of photoreceptor cells, and there are three in control flies). The expression of hRNH1 led to a significant increase of *ey>SmnRNAi* flies with normal eyes and heads (from 7% to 24%; Figure 3D), rescued the eye size and surface phenotypes, lowered the occurrence of reduced/rough eyes and decreased the frequencies of animals bearing abnormal ocelli (from 75% to 24%) or antenna alterations (from 20% to 5%; Figures 3F-3I). These findings collectively indicate that hRNH1 ectopic expression partially alleviates the eye development defects induced by Smn depletion, suggesting that an excess of R-loops reduces both the viability and the proliferative capacity of retinal precursors cells.

### Smn depletion causes alterations in transcriptome profiles that are partially rescued by hRNH1

To get insight into the molecular bases of the *Smn*-depletion phenotypes, we performed Illumina sequencing of RNA from whole *SmnRNAi* and control third instar larvae (minimum 150M paired-end reads/sample; principal component analysis (PCA) on transcriptomic profile showed clear separation between *Smn*-depleted and control samples (Figure S4A and Tables S1A-E). 31,050 annotated transcripts were quantified, with a minimum average Transcripts Per Million (TPM)>0.1 in at least one condition (75% of total reference transcripts). 1,015 genes were found to be differentially expressed between control and *SmnRNAi* larvae (|log2[Fold Change]|≥1 and adjusted p-value<0.05; Figure 4A and Table S1D). Gene ontology (GO) analysis (FDR<0.05) showed that the differentially expressed genes (DEGs) were enriched in nucleosome assembly and chromosome organization functions (supported by downregulation of replication-dependent histone isoforms; table S1D); other DEGs were involved in protein folding/proteolysis, response to heat and defense response (Figures 4B and 4C).

**Figure 4.**
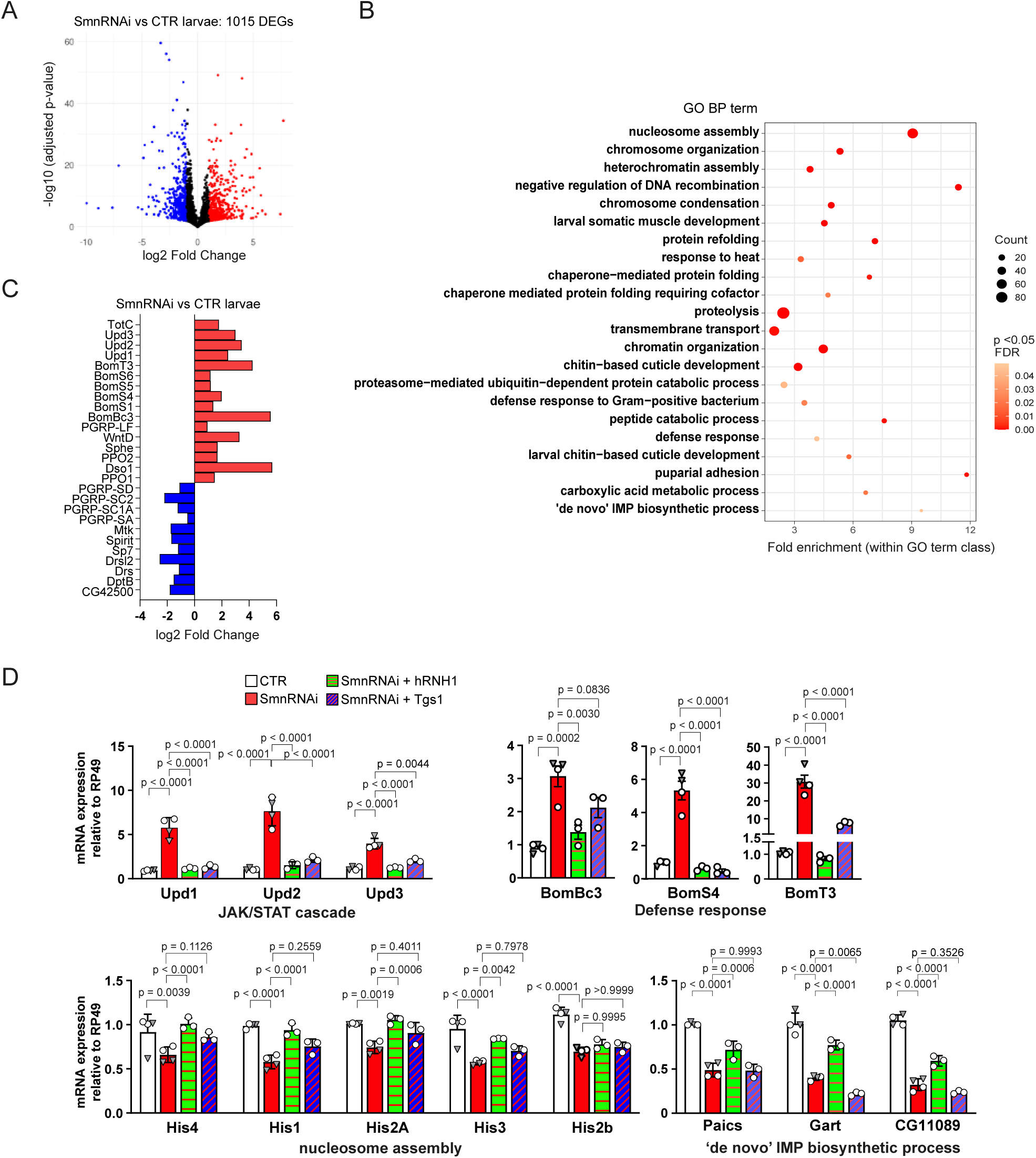
**Differential gene expression in *SmnRNAi, SmnRNAi + hRNH1 and SmnRNAi + Tgs1* larvae** (A) Volcano plot of differentially expressed genes (DEGs) in *SmnRNAi* vs control third instar larvae (blue and red dots: ǀlog2 [Fold Change] ǀ ≥1 and adjusted p-value <0.05). (B) Dot plot showing the GO Biological Process (BP) terms enriched within DEGs in *SmnRNAi* larvae. The size of each bubble reflects the number of DEGs associated with the GO term, while the color indicates the false discovery rate (FDR<0.05; Fisher’s exact test). (C) Bar plot showing the log2 [Fold Change] values of representative genes differentially expressed in *SmnRNAi* vs control larvae related to enriched GO terms for stress response and immune response. Bar plot showing log2 [Fold Change] values of representative genes differentially expressed in *SmnRNAi* vs. control larvae, related to enriched GO terms for stress and immune responses. (D) RT-qPCR validation of selected differentially expressed genes in *SmnRNAi* third instar larvae, compared to control, *SmnRNAi* + *hRNH1* and *SmnRNAi* + *Tgs1* larvae. Bars represent transcript expression levels (mean with SD) from at least three biological replicates (average of three technical replicates). For CTR and *SmnRNAi* larvae: 4 biological replicates include two RNA samples that were subjected to RNA-seq (white circles) or not (grey triangles). Data are normalized to *Rp49* and are relative to control larvae (set to 1). p values: two way-ANOVA with Tukey’s multiple comparisons test, genotype effect: F (3, 30) = 121.7, p <0.0001 (JAK-STAT cascade); F (3, 30) = 255.5, p<0.0001 (‘de novo’ IMP biosynthetic process); F (3, 50) = 59.86, p<0.0001 (nucleosome assembly); one way-ANOVA, F (3, 10) = 16.90, p=0.0003 (*BomBc3*); F (3, 10) = 51.83, p<0.0001 (*BomS4*); F (3, 10) = 47.88, p<0.0001 (*BomT3*).

The set of genes differentially expressed in *SmnRNAi* larvae was compared to DEGs from a previous study on *Smn^V72G^* early pupae that express a conserved variant residue, found in SMA patients ^30, 66^. We found 245 genes, that were differentially expressed in both mutant datasets, including genes related to the defense response and antimicrobial peptides (AMPs; Figures S4B and S4C). Among these, 193 genes were either up- or down-regulated in both conditions, with several involved in the innate immune response (Figure S4C and D). These include *PPO1* and *PPO2*, which are part of the melanization pathway, and the unpaired family cytokines *upd1, upd2*, and *upd3*, which activate the JAK/STAT pathway, triggering a systemic immune response ^67^. RT–qPCR analysis confirmed the upregulation of the three *upd* genes in *SmnRNAi* larvae compared not only to control but also to *SmnRNAi + Tgs1* and *SmnRNAi + hRNH1* larvae (Figure 4D). The activation of the innate immune response is mediated by the conserved Toll and Immune-deficiency (Imd) pathways that stimulate the production of AMPs effectors ^68^. Six out of the twelve members of the Bomanin family, the principal effectors of the Toll pathway ^69, 70^, showed increased expression in *SmnRNAi* larvae, compared to control (*BomBc3, BomS1, BomS4, BomS5, BomS6, BomT3*; Table S1D); RT–qPCR analysis for *BomBc3, BomS4* and *BomT3*, confirmed their upregulation. Importantly, the transcription levels of these 3 genes returned to baseline in *SmnRNAi + hRNH1* larvae, while *BomS4* and *BomT3* levels were decreased in *SmnRNAi + Tgs1* larvae (Figure 4D). RT-qPCR also confirmed the downregulation of several histone isoforms in *SmnRNAi* larvae, compared to control larvae (Figure 4D and Table S1D). *SmnRNAi + hRNH1* larvae, but not *SmnRNAi + Tgs1* larvae, showed normal expression of mRNAs encoding *His1, His2A, His3, His4*; while the *His2B* mRNA level was not rescued in either *SmnRNAi+hRNH1* or *SmnRNAi+Tgs1* larvae (Figure 4D). Interestingly, also genes implicated in the de novo purine biosynthesis pathway such as *Paics*, *Gart* and *CG11089/ATIC* showed downregulation in *SmnRNAi* and in *SmnRNAi* +*Tgs1* larvae, whereas their levels were normal in *SmnRNAi + hRNH1* larvae (Figure 4D).

### hRNH1 and Tgs1 impact the gene expression and splicing profiles in SmnRNAi brains

To further explore how hRNH1 and Tgs1 affect the *Smn* loss of function phenotypes in the nervous system, we performed Illumina sequencing (minimum 120M paired-end reads/sample) to characterize the total transcripts in brains from *SmnRNAi, SmnRNAi + Tgs1, SmnRNAi + hRNH1* larvae and the corresponding controls (Table S2A-H); gene expression-based sample clustering clearly recapitulated sample conditions (Figure S5A). We detected 31,025 known transcripts (TPM>0.1 across replicates in at least one condition), which correspond to 10,492 genes. 272 genes exhibited differential expression between *SmnRNAi* and control brains (|log2[Fold Change]|≥1 and adjusted p-value <0.05; Figure S5B and Table S2E). GO term enrichment analysis (Figure 5A) identified top biological processes related to nucleosome assembly, proteolysis, protein folding and response to heat stress. We also found deregulation of snRNAs and proteins related to DNA damage response. Comparison of the differential expression analyses performed on brain and larval RNA datasets, relative to their respective controls (Tables S2E and S1D), showed 136 genes, whose expression was altered in both datasets, including *upd1-3* and several histones (Figure S5C).

**Figure 5.**
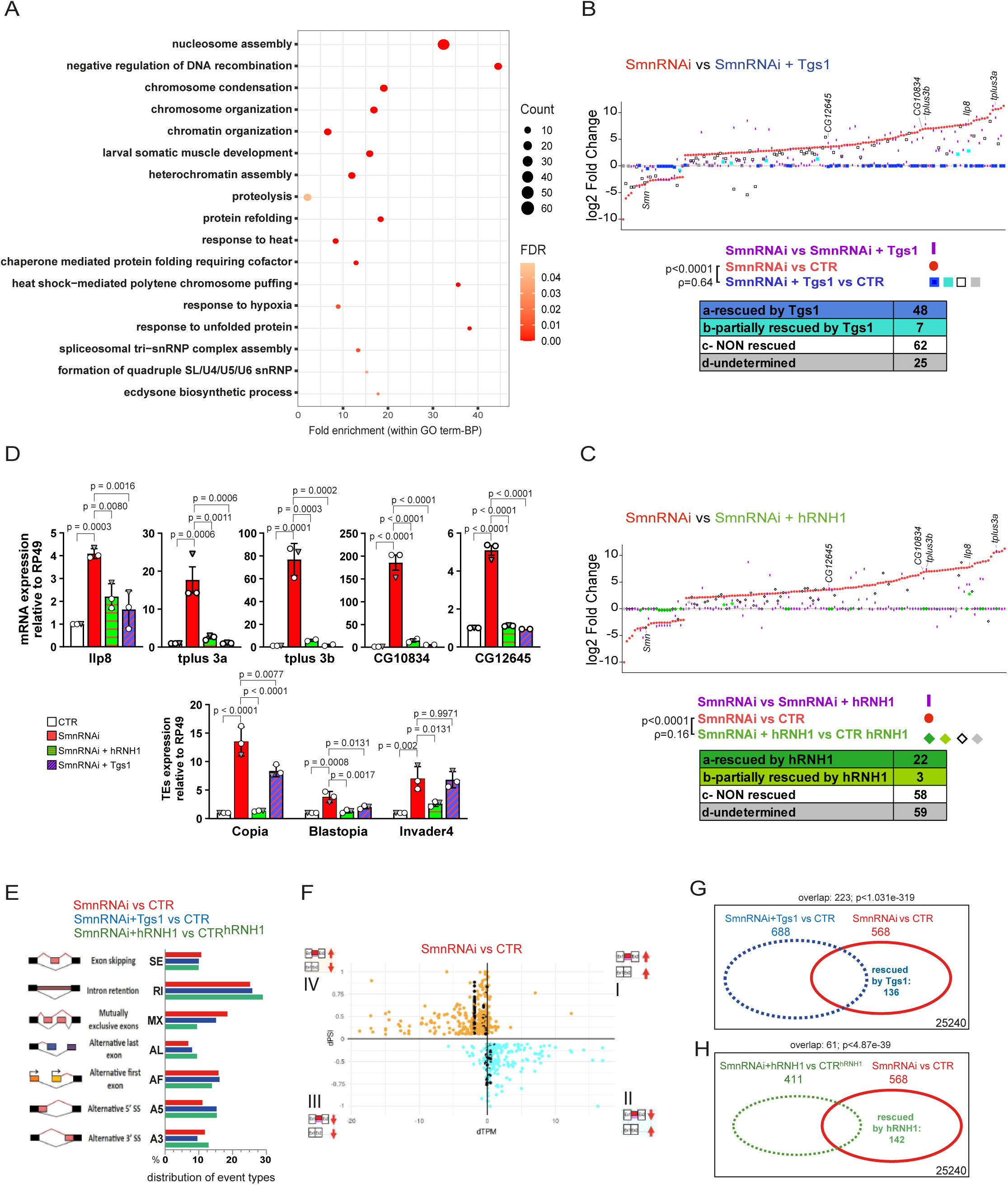
**Altered gene expression and splicing profiles in *SmnRNAi* brains are partially rescued by hRNH1 or Tgs1** (A) Bubble plot showing the GO Biological Process (BP) terms enriched within DEGs in *SmnRNAi* vs control brains. The size of each bubble reflects the number of DEGs associated with the GO term, while the color indicates the false discovery rate (FDR<0.05; Fisher’s exact test). (B, C) Representation of 142 significantly varying DEGs in *SmnRNAi* vs CTR brains (|log2[Fold Change]ǀ ≥2 and adjusted p-value<0.05). Colored symbols: log2[Fold Change] of DEGs in pairwise comparisons between the indicated conditions: (i) *SmnRNAi vs*. control (red dots); (ii) *SmnRNAi* + rescue construct *vs*. control (*Tgs1*, squares in B; *hRNH1*, diamonds in C); (iii) Smn*RNAi vs. SmnRNAi +* rescue construct (purple dashes). Significance and correlation of the pairwise comparisons (i) and (ii) were assessed using the two-tailed Wilcoxon matched-pairs signed rank test (p-value) and two-tailed Spearman’s correlation (ρ). Rescue effects are ranked from *a* to *d*; symbols labeling condition ii (*SmnRNAi* + rescue construct *vs*. control) in graphs B and C, are color-coded according to their class. See also Figure S6 and Table S2G. (D) RT–qPCR analysis of the expression levels of selected DEGs and transposable elements (TEs) in brains with the indicated genotypes. Data from three biological replicates (average of three technical replicates per sample) are shown relative to *Rp49* and normalized to control samples. Bars: mean, SD; p-values: one-way ANOVA with Tukey’s multiple comparisons test; F (3, 8) = 20.79, p=0.0004 (*Ilp8*); F (3, 6) = 58.90, p<0.0001 (*tplus3b*); F (3, 8) = 22.22, p=0,0003 (*tplus3a*); F (3, 5) = 62.42; p=0.0002 *(CG10834);* F (3, 7) = 208.5, p<0.0001 *(CG12645);* F (3, 8) = 57.94, p<0.0001 *(Copia)*; F (3, 8) = 17.45, p =0,0007 *(Blastopia)*; F (3, 8) = 16.48, p=0,0009 *(Invader-4)*. (E) Distribution of differential alternative splicing (AS) events among seven major classes in the indicated pairwise comparisons, │dPSI│≥0.3; adjusted p-value<0.05. (F) Partitioning of 568 significant AS events in *SmnRNAi* brains (│dPSI│≥0.3 and adjusted p-value<0.05) based on dPSI and dTPM values. Quadrants IV and II contain AS events, where the expression levels of the isoform lacking the AS inclusion event in *SmnRNAi* brains are lower (quadrant IV), and higher (quadrant II), compared to controls. Specifically, quadrant IV contains upregulated “AS inclusion isoforms” with downregulation of the exclusion isoform, while quadrant II contains events with the opposite behavior. Black dots represent 103 AS events mapping to *Dscam1* transcripts. dPSI indicate the proportion of reads with the inclusion of AS events relative to inclusion and exclusion reads (see also Figure S7A). (G, H) Venn diagrams representing the number of overlapping differential AS events between comparisons. 25,240: total events; 136 and 142 AS events showed improvement, in *SmnRNAi + Tgs1* and *SmnRNAi + hRNH1* brains, respectively. p-values were calculated *via* hypergeometric test. See also Figures S7C,F and Table S3B).

To evaluate whether the expression of Tgs1 and hRNH1 impacts the expression of DEGs in *SmnRNAi* brains, we focused on a subset of 142 DEGs in brain datasets, with |log2[Fold Change]|≥2 (Table S2E) and analyzed the relative changes in the expression levels of each gene by performing pairwise comparisons between genotypes, as outlined in Figures 5B, 5C, S6 and Table S2G. The efficiency of *Smn* knockdown was uniform across the three *Smn-*depleted datasets, as measured by RT-qPCR (Figure S5D). For each comparison, we assigned the DEGs to four classes, ranging from fully rescued to not rescued (class a to d; Figure S6). We defined a DEG as “rescued” by either Tgs1 or hRNH1 if it was differentially expressed in the comparison “*SmnRNAi vs CTR”* but not in the comparison “*SmnRNAi + rescue construct* (*Tgs1* or *hRNH1*). Overall, we observed a substantial reduction in the |log2[Fold Change]| values when comparing *SmnRNAi* samples without and with the expression of either rescue construct (Figures 5B, 5C, S6 and Table S2G). A subset of differentially expressed genes rescued by both hRNH1 and Tgs1 (*CG12645, CG10834, Ilp8, tplus3a, tplus3b*) was validated by RT-qPCR (Figure 5D).

To improve transcript reconstruction and to characterize unannotated transcript isoforms resulting from defective splicing, we performed long-read Oxford Nanopore Technology (ONT) sequencing on *SmnRNAi* mutant brains and controls. Combining Illumina and ONT approaches enables the detection of multiple altered splicing events occurring within the same isoform ^10^. At least 1.3M ONT multiexonic reads per sample were generated (Table S2B), mapped to the fly genome and used for transcript assembly and quantification with FLAIR ^71^, with reconstructed isoforms supported by at least 3 ONT reads. The “extended transcriptome” we generated included 43,715 transcripts (of which 18,434 were annotated and 25,281 were novel transcripts), mapping to 20,950 loci (Table S2F). Among these, 4,331 derived from repetitive element insertions (Table S2F). Transposable elements expression was assessed both on brain and whole larval RNA datasets (see methods; Tables S1E and S2H). The expression levels of *Copia*, *Blastopia* and *Invader-4* retrotransposons, whose expression increases with aging ^72, 73, 74^, were upregulated in both *SmnRNAi* brains and whole larvae compared to controls (Figure S5E) and were restored to normal levels in brains upon expression of hRNH1; *Copia* and *Blastopia* levels were reduced upon expression of Tgs1 (Figure 5D). The extended transcriptomes were used to explore splicing profiles. We used SUPPA2 ^75^ for detection of differential alternative splicing (AS) events. 25,240 events were detected and classified into seven major AS event types (Figures 5E, S7A and table S3A). In Figure 5F, AS events are distributed across four quadrants (I-IV) based on their Delta Percent-Spliced-In (dPSI) values, plotted against the expression levels (TPM) of the corresponding isoform without the AS event. To exclude AS events that are merely the consequence of varied transcriptional activity at the corresponding locus, we focused only on events in quadrants II and IV, where the expression levels of the isoform with and without the AS event were discordant (about 80% of the total events; Figure S7B). To evaluate potentially rescued events, we focused only on significant AS events, in *SmnRNAi* brains, with │dPSI│≥ 0.3. This filtering resulted in 568 AS events in the *“Smn RNAi vs CTR”* comparison; 688 AS events in “*Smn RNAi + Tgs1 vs CTR”* and 411 in “*Smn RNAi + hRNH1 vs CTR-hRNH1”* (Figures 5G and 5H). Next, we focused on the 568 significant *SmnRNAi* AS events, whose distribution into AS classes is reported in Figure 5E and evaluated the status of the event in the other two conditions through pairwise comparisons between conditions (Figures 5G, 5H and Table S3B). We explored potentially rescued events as outlined in Figures S7C and S7D. Of the 568 significant AS events altered in *SmnRNAi brains*, 136 and 142 showed rescue in *SmnRNAi + Tgs1* and *SmnRNAi + hRNH1* brains, respectively (Figures 5G, 5H and S7E,F). Finally, to confirm the robustness of the SUPPA2 analyses, we validated an intron retention event in the *Med6* gene through RT-qPCR analysis. This event had a dPSI of 0.17, falling below the │dPSI│≥0.3 threshold set as relevant for our analysis (Figures S8A-C).

Overall, AS events detected in *SmnRNAi* brains did not result in a net significant change in the expression levels of the corresponding genes (Table S3C). While AS events could affect the expression levels of specific protein isoforms, splicing deficiency per se appears to be insufficient to fully explain SMA pathogenesis, which depends also on other *Smn-*controlled processes, such as the delivery of specific transcripts to ribosomes, which may play a significant role in the differential expression of proteins and the proteostatic deficits seen in many SMA models ^23, 24, 30, 76, 77^. In summary, our findings demonstrate that splicing defects have a significant impact on the epigenome, potentially leading to transcriptional stress and the formation of R-loops, which could in turn further affect splicing efficiency. These events, coupled with the histones deficit, could ultimately result in DNA damage, which would contribute to neurosenescence and activate the innate immune response, leading to systemic inflammation.

### Smn depletion induces glutamate, aspartate, and serine alterations, restored by hRNH1

High-Pressure Liquid Chromatography (HPLC) analysis on the CSF of pediatric SMA patients and on the central nervous system of SMNΔ7 mice ^37, 38, 39^ indicated a prominent influence of SMN in regulating the levels of neuroactive amino acids acting on glutamatergic neurotransmission. Here, we investigated whether Smn depletion in *SmnRNAi* larvae similarly affects the concentration of D- and L-amino acids modulating glutamate receptor activation (Figures 6A-M). As observed also in severe SMA patients ^39^, HPLC measurements in *SmnRNAi* larvae highlighted a significant decrease in L-Glu concentration ratio along with an upregulation of L-Gln/L-Glu ratio compared to control larvae (Figures 6B–Figures 6BD). Moreover, further suggesting an abnormal Glu metabolism in *SmnRNAi* larvae, we also report a significant increase in the GABA/L-Glu ratio (Figure 6F). Besides L-Glu variations, in *SmnRNAi* larvae, we determined a striking downregulation of L-aspartate (Figure 6I), along with a significant accumulation of its D-enantiomer (Figure 6H) and enhanced D-Asp/total Asp ratio (Figure 6J). Notably, we found a significant reduction of the D-Ser/total Ser ratio in *SmnRNAi* larvae (Figure 6M), overall indicating an unexpected influence of Smn levels in modulating aspartate and serine enantio-conversion. Taken together, our observations in *SmnRNAi* larvae well align with the abnormally low concentrations of glutamate coupled to higher values of Gln/Glu ratio reported in the CSF of SMA type 1 patients and in the spinal cord and brain of SMNΔ7 mice, confirming a prominent role of Smn in modulating amino acids regulating glutamate receptors activation.

**Figure 6.**
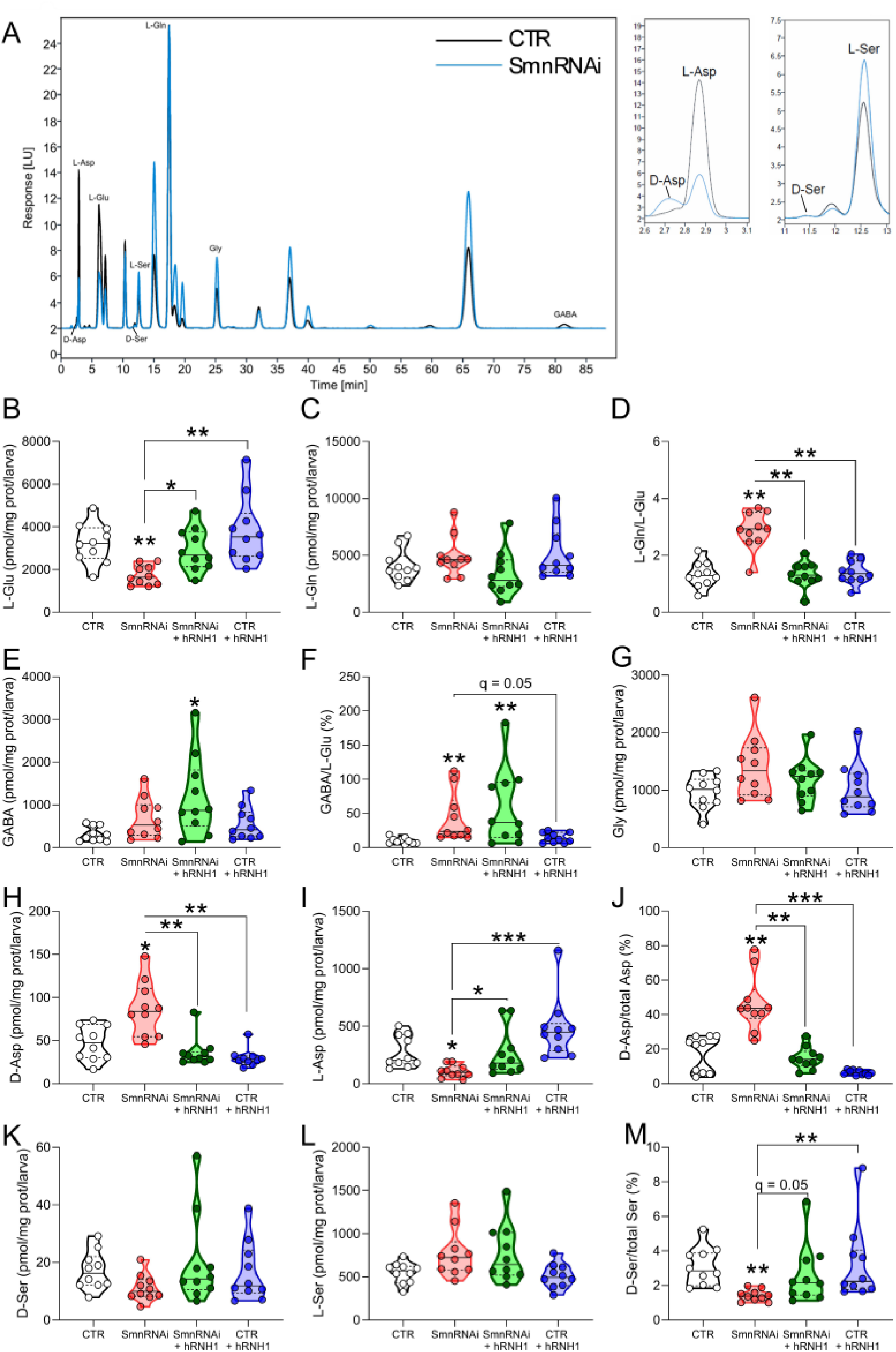
**Smn depletion induces glutamate, aspartate, and serine metabolism alterations that are restored by hRNH1** (A) Representative chromatogram showing the peaks of D-aspartate (D-Asp), L-aspartate (L-Asp), L-glutamate (L-Glu), D-serine (D-Ser), L-serine (L-Ser), L-glutamine (L-Gln), glycine (Gly), and γ-aminobutyric acid (GABA) in whole CTR and *SmnRNAi* larvae, with magnification of D-Asp and L-Asp as well as D-Ser and L-Ser peaks. (B-M) Levels of L-Glu (B), L-Gln (C), L-Gln/L-Glu percentage ratio (D), GABA (E), GABA/L-Glu percentage ratio (F), Gly (G) D-Asp (H), L-Asp (I), D-Asp/total Asp percentage ratio (J), D-Ser (K), L-Ser (L) and D-Ser/total Ser percentage ratio (M) in the indicated groups of CTR, *SmnRNAi*, *SmnRNAi + hRNH1*, *CTR + hRNH1* larvae. Data are shown as violin plots representing the median with interquartile range (IQR). \**q*<0.05, \*\**q*<0.01, \*\*\**q*<0.0001 (non-parametric Kruskall-Wallis test followed by original false discovery rate method of Benjamini and Hochberg’s *post-hoc* for multiple comparisons). Dots represent values from each sample analyzed, n = 10/group.

Then, to explore the impact of R-loops on amino acid abnormalities, we measured the content of these biomolecules in *SmnRNAi + hRNH1* larvae. Noteworthy, in comparison to *SmnRNAi* larvae, the *SmnRNAi + hRNH1* larvae showed higher L-Glu (Figure 6B) and L-Asp (Figure 6I) concentrations coupled with increased D-Ser/total Ser ratio (Figure 6M). Conversely, when we compared *SmnRNAi* to *SmnRNAi + hRNH1* larvae, we found a significant decrease in D-Asp concentrations (Figure 6H) along with lower L-Gln/L-Glu (Figure 6D) and D-Asp/total Asp (Figure 6J) ratios. Thus, HPLC determination highlighted a rescue effect of hRNH1, leading to comparable D- and L-amino acid concentrations between *SmnRNAi + hRNH1* and control larvae. In contrast, the GABA/L-Glu ratio was not rescued by hRNH1 expression in *SmnRNAi* larvae (Figure 6F). Finally, hRNH1 expression in control larvae did not perturb the D and L amino acid content (Figure 6B-M).

Interestingly, the reported amino acids changes in *SmnRNAi* larvae were associated with alterations in the expression levels of genes encoding enzymes involved in their metabolism (listed in Figure S9A). Among the most significantly dysregulated genes found in SmnRNAi larvae, some are implicated in clinically relevant pathways ^37, 38, 39^ such as catecholamine (*Dopa decarboxylase, Ddc*), glutamate (glutamine synthetase, *Gs1* and *Gs2*), and glycine metabolism (*serine hydroxymethyltransferase, Shmt*). Their downregulation was confirmed by RT-qPCR in both *SmnRNAi* and *SmnRNAi + RNH1* larvae (Figure S9B). These results are consistent with studies performed in mice ^19^, suggesting that these enzymatic changes are a response to low Smn levels. *Ddc* downregulation was also observed in SMA derived cells and overt symptomatic SMA mouse models^38^, thus reinforcing the translational relevance of the fly model in recapitulating mechanisms pertinent to SMA.

In conclusion, our results indicate that changes in glutamate, aspartate, and serine metabolism in larvae are closely linked to the accumulation of R-loops induced by Smn downregulation. However, it remains still unclear whether these amino acid differences can be regarded as indicators of varying cell composition in the *SmnRNAi* larval brains or whether they are associated with disruptions in the cell cycle, inflammation, altered metabolism or cell death.

## DISCUSSION

The mechanisms linking SMN deficiency to motor neuron loss remain elusive. Among the plethora of SMN functions, recent evidence suggests that SMN may also prevent the accumulation of RNA:DNA hybrids, generating R-loops, whose excessive presence can result in DNA damage, immune response and disease^44, 50, 52, 63, 78, 79^. Here we have shown that Smn depletion in *Drosophila* augments R-loop formation, just like the deficiency of its human ortholog ^44, 52, 78^, leading to DNA damage and its detrimental cellular responses.

The central finding of our study is that prominent genetic, cellular and biochemical abnormalities that we observe in Smn-deficient flies are largely suppressed by the overexpression of hRNH1, highlighting the impact of transcriptional stress and DNA damage on Smn-dysfunction (Figure 7). Although ectopic hRNH1 expression does not restore the *Smn* transcript level, or rescues the lethality of *SmnRNAi* flies, it does suppress

**Figure 7.**
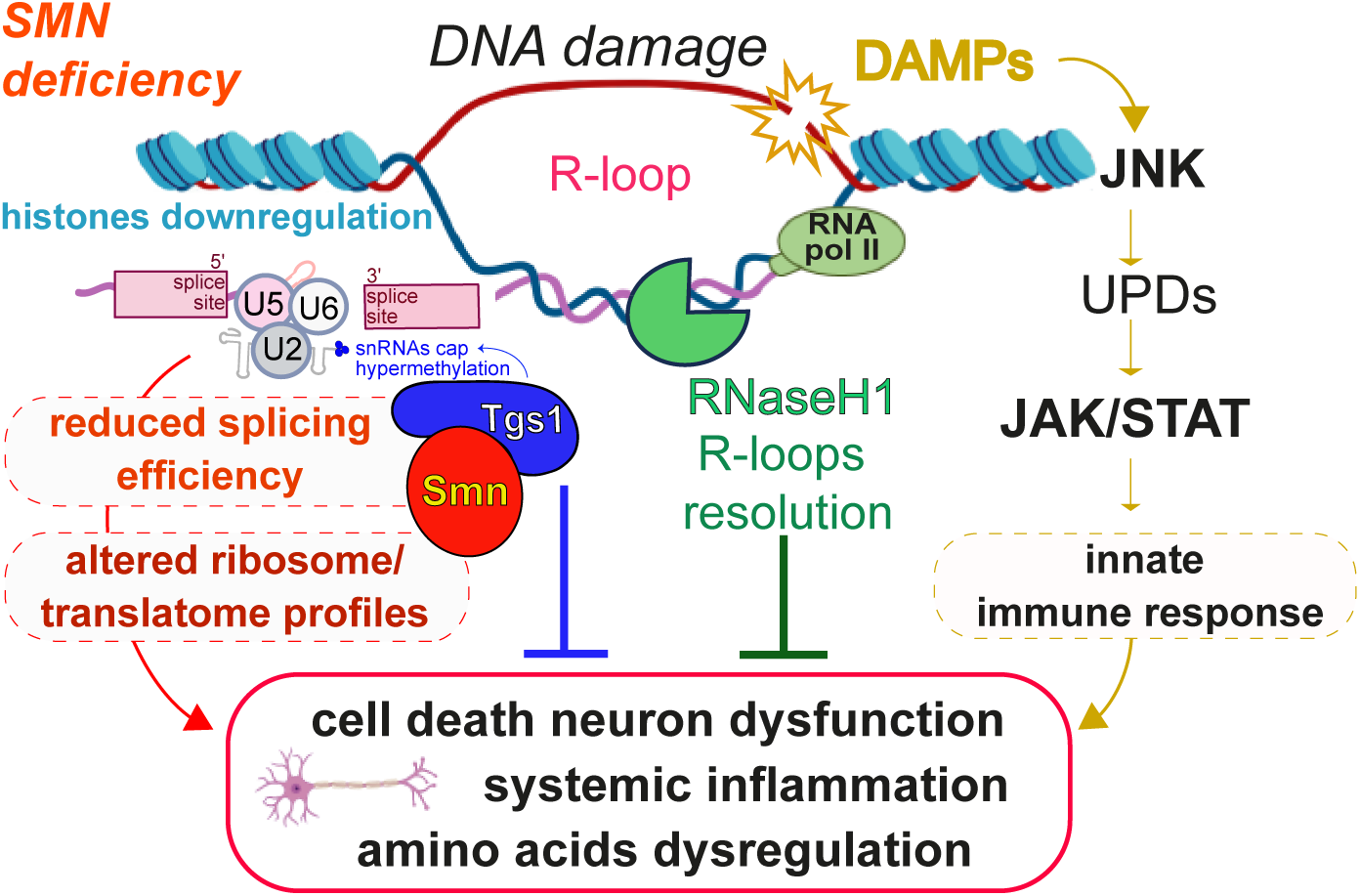
Model illustrating the consequences of Smn dysfunction, highlighting the role of disrupted splicing and RNA:DNA hybrids in the activation of the innate immune response, mitigated by the expression of hRNH1 or the overexpression of Tgs1. See text for a detailed explanation of the model.

R-loop-dependent DNA damage and its consequences. In addition, transcriptome analyses revealed that hRNH1 normalizes the expression of a subset of genes deregulated in response to Smn deficiency and improves splicing efficiency.

The mRNA alterations due to splicing defects are insufficient to fully explain both SMA pathogenesis ^24, 77^ and the Smn-related *Drosophila* phenotypes^76^. Indeed, we argue that the splicing defects are likely to promote excessive R-loop formation, which could contribute to most of phenotypic defects caused by Smn deficiency (Figure 7). Specifically, in brains from *SmnRNAi* larvae, we have detected DNA damage and shown that it can result in cell cycle delays and an accelerated differentiation of ganglion mother cells toward the neural fate, possibly at the expense of gliogenesis. Premature differentiation of neuroblasts has been described as a response to various genotoxic stressors, including irradiation, oxidative damage, or aneuploidy ^80, 81, 82, 83^. DNA damage in *SmnRNAi* larval brains might therefore alter the neuron-to-glia shift in progenitor differentiation, which occurs during larval neurogenesis ^84, 85^. Importantly, hRNH1 expression in *SmnRNAi* brains restored both cell cycle dynamics and the balance between glia and neurons. In addition, hRNH1 expression in the eye imaginal discs reduced both the decay of photoreceptor precursors in the eye and the head defects caused by Smn deficiency.

Collectively, these results show that Smn depletion affects neural cell viability and differentiation and indicate that these effects result from an R-loop induced chronic DNA damage.

In line with previous results^30^, we confirmed that *Smn* deficiency leads to innate immune response activation, causing an upregulation of antimicrobial peptides and formation of melanotic masses. Notably, the increased levels of pro-inflammatory cytokines Upd 1-3 and Bomanins in *SmnRNAi* larvae and the formation of melanotic masses were both suppressed by ectopic hRNH1 expression. This finding suggests that the RNA:DNA hybrids and single-stranded DNA exposure in R-loops, are perceived as damage-associated molecular patterns (DAMPs) and activate the p53-mediated JNK pathway, leading to cytokine upregulation and initiation of the JAK-STAT signaling pathway ^67, 86, 87, 88^ (Figure 7).

Furthermore, consistent with the fact that Tgs1 interacts both physically and functionally with Smn^7^, we observed that it rescues many, but not all the abnormalities that are rescued by hRNH1. Indeed, overexpression of either Tgs1 or hRNH1 rescued the cell cycle defects and mitigated cytokine upregulation and inflammatory responses, underscoring the links between Smn, Tgs1, and RNH1 in R-loop metabolism and inflammation.

Interestingly, we also found that hRNH1 expression counteracts the amino acid homeostasis disruption induced by Smn depletion. In *SmnRNAi* larvae, the variations in glutamate, serine and aspartate, which play essential roles for energy homeostasis-related processes, purine and pyrimidine metabolism, neuroinflammation ^89, 90, 91^, and neurotransmission ^37, 38, 39, 40, 58, 92, 93^, were significantly rescued by reducing RNA:DNA hybrids. However, the molecular mechanism leading to these variations remain to be elucidated.

Overall, the relationship between R-loops and immunological responses reveals a cellular crosstalk that is likely to underlie the multiple phenotypes observed in SMA patients and models. The accumulation of DNA damage due to Smn deficiency affects the behavior of neuronal lineages, potentially inducing senescent phenotypes that alter cell fate specification. Thus, given the emerging role of immune modulation in alleviating neuroinflammation and neurodegeneration associated with SMA ^27^, it is conceivable that targeting the immune system could relieve symptoms in SMA patients. hRNH1 expression or R-loop reduction may also offer a way to partially mitigate the consequences of Smn-related systemic inflammation.

In *SmnRNAi* larvae, DNA damage suppression by hRNH1 or Tgs1 overexpression normalizes the expression levels of several genes, including the histones genes and the retrotransposons, and leads to a reduction of aberrant splicing isoforms, highlighting the role of chromatin remodeling in regulating splicing efficiency and transcription accuracy. Smn likely plays both direct and indirect roles in these processes, possibly controlling snRNP quality and interacting with RNA polymerase II ^41, 94^. This interaction may create an epigenetic environment for effective mRNA co-transcriptional processing, termination, export, and ribosome recognition, ultimately enhancing the translation efficiency of mRNAs.

Although it remains still unclear how DNA damage affects postmitotic neurons, metabolic pathways, neurotransmission and inflammatory response in SMA patients and mammalian models ^27, 95^, the beneficial effects of hRNH1 in alleviating genotoxic stress in *Drosophila* pave a new avenue for further studies aimed at improving SMA outcomes.

By linking the immunomodulatory role to reduced DNA damage, R-loop modulation by hRNH1 or other treatments, could potentially extend the benefits of SMN replacement therapies, to ameliorate genotoxic components, particularly in peripheral tissues where correction is less effective. We hypothesize that alleviating the immunomodulatory consequences of chronic transcriptional stress could enhance the efficacy of SMN-restoring therapies in the nervous system, particularly in cells where significant SMN level increases are unattainable.

## Materials and Methods

### *Drosophila* strains and transgenic constructs

The *UAS-Smn RNAi* construct [P/TRiP.HMC03832/attP40 (UAS-*Smn*RNAi)] and the *Actin-GAL4* driver were obtained from the Bloomington Stock Center. The deficiency that removes the *Smn* gene (the *Smn^X7^*allele) is a gift from Dr. Artavanis-Tsakonas. The *UAS-GFP-dTgs1* strain carries the pPGW*-Tgs1*construct, generated by cloning the *dTgs1* CDS into the pPGW destination vector (stock number 1077, Drosophila Genomics Resource Center, supported by NIH grant 2P40OD010949), using Gateway technology (Thermo Fisher Scientific). The UAS-hRNH1 line^62^ was kindly provided by Dr Fernando Azorín. For the generation of WT and catalytically inactive RNaseH1-GFP constructs, we subcloned into the pJZ4 plasmid the RNaseH1 coding sequences from the constructs reported in^96^ (kindly provided by Dr Jana Dobrovolná), flanked by KpnI and Xbal sites. The Oregon-R strain was used as a wild-type control. All flies were reared according to standard procedures at 25°C. Lethal mutations were balanced over either *TM6B, Hu*, *Tb* or *CyO-TbA, Cy, Tb* chromosome balancers.

### NMR sample preparation, spectra acquisition and statistical analysis

^1^H-NMR metabolomics experiments were carried out on 50 third instar larvae. The extraction was carried out in a double phase, placing the larvae into a mixture of H_2_O, CH_3_OH, and CHCl3 equal to 48, 240, and 200 µl, respectively. The samples were sonicated for 2 minutes using an ultrasonic tip sonicator (Vibracell VCX 750,Sonics®). 120 μL of H_2_O and 120 μL of CHCl_3_ were added to the mixture. The polar phase was taken after centrifugation at 10.000 g for 15 minutes. The solvents are removed using SP Genevac EZ-2 4.0 Series Centrifugal Evaporators. Dried polar extracts were dissolved into 200 μL of buffer (50 mM Na_2_HPO_4_, 1 mM trimethylsilyl propionic-2,2,3,3-d4 acid, sodium salt (TSP-d_4_), 50 μL of D_2_O) and transferred into 3 mm NMR tubes for ^1^H NMR detection. TSP-d_4_ at 0.1% in D_2_O was used as an internal reference for the alignment and quantification of the NMR signal. A Bruker DRX600 MHz spectrometer (Bruker, Karlsruhe, Germany) fitted with a 5 mm triple-resonance z-gradient TXI Probe was used for NMR investigations. 1D-NOESY experiments were acquired at 298 K with the excitation sculpting pulse sequence to suppress the water resonance at a 12 ppm sweep width, 128 transients of 19 k complex points, with acquisition time of 4 sec transient and 10 msec mixing time. The pre-processing phase was performed by applying the Fourier transform followed by the manual phase and baseline correction. TOPSPIN, version 3.2, was used for spectrometer control and data processing (Bruker Biospin, Fällanden, Switzerland). Spectra assignment was performed using Chenomix and quantified using NmrProcFlow, as previously reported.^97^

Partial least square determination analysis (PLS-DA) was performed on the normalized dataset using MetaboAnalyst 6.0 (http://www.metaboanalyst.ca/). The model resulting from the supervised approach was validated by 10-fold cross method considering the Q2, R2 indices and accuracy. ^98^ The contribution of the individual variables in generating the separation of the clusters was carried out using Variable Importance Projection (VIP), considering significant only the metabolites with VIP>1.2. The hierarchical clustering analysis was carried out using MetaboAnalyst 6.0, considering the median metabolomic profile^99^. Clustering was performed using Ward’s method, and the distance between the clusters was calculated based on the Euclidean distance.

The Pathway Topology analysis on the *Drosophila Melanogaster* database was performed using MetPa^100^ . Pathways with a number of Hits, i.e. metabolites belonging to the biochemical pathway>2 and p-value<0.05, were considered significant. To understand the influence of individual pathways on the examined clusters, the Pathway Impact (PI) value was considered^100^. The PI was calculated by combining the path centrality and enrichment results.

### Dot Blot analysis

Dot blot analysis was performed as previously described ^112^. Briefly, ∼40 whole third instar larvae were pestled in liquid nitrogen with mortar, powder was resuspended in 1ml lysis buffer (50 mM Tris-HCl pH 8.0, 100 mM EDTA,100 mM NaCl 0.5% SDS) adding 500ng/ul proteinase K. The reaction was incubated overnight at 37°C. Genomic DNA was then purified with phenol/chloroform and treated with 5 units of RNase III (Thermo Fisher Scientific AM2290 Ambion™) either alone or in combination with 25 units of RNase H1 (EN0201 Thermo Scientific™) and incubated for 2h at 37°C. After phenol/chlorofom purifiation, 1:2 serial dilution of genomic DNA (starting from 500ng) was spotted on Hybond-N+ hybridization membrane, cross-linked to the membrane by UV ligh (1,200uJx100) and incubated overnight with anti-S9.6 1:1.000 (MABE, MABE1095 1095 anti-DNA-RNA Hybrid, clone S9.6) and anti-dsDNA 1:10.000 as loading control (Abcam ab27156 Anti-dsDNA antibody).

### Immunostaining

Third instar larval brains were dissected in PBS and fixed in 1,8% formaldehyde and 45% acetic acid for 10 min at room temperature (for RNA:DNA hybrids detection), or fixed in 3,7% formaldehyde for 20 minutes and in 45% acetic acid for 2 minutes at room temperature (for ɤH2Av, pH3, ELAV and Repo detection). The brains were then squashed and the slides were placed for at least 5 minutes in liquid nitrogen. After removal of the coverslip, squashed brains were washed in PBS 0.1% Triton X-100 (PBS-T) for 2 x 10 min and blocked with PBS-T and 5% BSA for 60 min. For immunostaining the brains were incubated overnight at 4 °C with mouse anti-S9.6 (1:300; Merk Millipore), mouse anti-ɤH2Av (1:20;DSHB), rabbit anti-phospho-Histone3(Ser10) (1:250; EMD Millipore Corporation), mouse anti-ELAV (1:50; DSHB) or mouse anti-Repo (1:100; (DSHB). The slide were then washed 2 x 10 min in PBS and then incubated for 1 h at room temperature with Cy3-conjugated anti-mouse for the mouse S9.6 and ɤH2Av (1:50, Jackson Immunoresearch), FITC-conjugated anti-mouse for the mouse ELAV and Repo (1:20, Jackson Laboratories), or with Cy3-conjugated anti-rabbit for the rabbit pH3 (1:300, Jackson Laboratories). All samples were mounted in Vectashield with DAPI (Vector) to stain DNA. S9.6, ɤH2Av, pH3, ELAV and Repo positive foci were quantified using Zen 2.5 Pro software (Zeiss, Germany). Image z-stacks were acquired with the same parameters using an Axiocam 512 (Zeiss) monochromatic camera. Fluorescence threshold was setup starting from the basal fluorescence, and spots were considered positive starting from 3 times the basal fluorescence. Statistical significance was calculated using GraphPad Prism Software 10.4.1

### EdU-labeling

Mid third-instar larval brains were dissected in 0.7% sodium chloride solution (NaCl) and incubated for 2 h at 25 °C in phosphate-buffered saline (PBS) with 100 μM 5-Ethynyl-2’-deoxyuridine (EdU). Brains were then fixed in 4% formaldehyde in PBS at room temperature for 20 min, and after a brief incubation in 45% acetic acid, they were squashed in 60% acetic acid and frozen in liquid nitrogen. After removal of the coverslip, the slides were immersed for 10 min in ethanol at –20 °C, washed two times in PBS + BSA 3% (5 min each), washed for 20 min in PBT (PBS containing 0.2 % Triton X-100), and then incubated for 30 minutes with the Click-iT reaction cocktail according to the Click-iT Edu Imaging Kits protocol (Invitrogen). All samples were mounted in Vectashield with DAPI (Vector) to stain DNA. Images acquisition and statical analysis were done as above.

### TUNEL assay

Mid third-instar larval brains were dissected in phosphate-buffered saline (PBS) and incubated in 4% paraformaldehyde in PBS at room temperature for 20 min. After fixation, the tissues were washed in PBS 0.2% Triton X-100 (PBS-T) for 3 x 10 min. Brains were incubated overnight at 4°C with mouse anti-ELAV [1:100; Developmental Studies Hybridoma Bank (DSHB)]. Brains were then washed 3 x 10 min in PBS 0,2% Triton X-100 (PBS-T) and incubated for 1 hr at room temperature with Cy3-conjugated anti-mouse (1:50, Jackson Laboratories). After further washing (3 x 10 min in PBS 0.2% Triton X-100), brains were incubated at 37 °C for 2 h in the TUNEL mixture in the dark according to the *In Situ* Cell Death Detection Kit protocol (Roche). Some brains were incubated only with the label solution as a negative control and, as expected, they gave no TUNEL signal. Brains were then washed in PBS-T for 3 x 10 min and mounted in Vectashield with DAPI (Vector) to stain DNA and reduce fluorescence fading. Images were acquired using a CrestOpticsV3 confocal spinning disk mounted on a Nikon Eclipse Ti2-E inverted microscope with a Last generation Perfect Focus System. Images represent the Maximum Intensity Projection (MIP) of 61.28 µm Z-stacks (z. step 0.30 µm). Tunel and ELAV positive nuclei were quantified using Fiji software.

### Microscopic analysis of Eye-antennal imaginal discs

Microscopic analysis of Eye-antennal imaginal discs Third instar larval eye-antennal imaginal discs were dissected in ice cold PBS and fixed in 4% paraformaldehyde in PBS at room temperature for 30 min. After fixation, the tissues were washed in PBS 0,3% Triton X-100 (PBS-T) for 3 x 20 min and blocked with PBS-T and 5% BSA for 30 min. Discs were incubated overnight at 4°C with anti-Cleaved Caspase3 (1:300; Cell Signaling Technology) and mouse anti-Elav [1:100; Developmental Studies Hybridoma Bank (DSHB)], washed 3 x 20 min in PBS-T, and then incubated for 1 hr at room temperature with Cy3-conjugated anti-rabbit (1:300, Life Technologies) and FITC-conjugated anti-mouse (1:100, Jackson – Laboratories). All samples were mounted in Vectashield with DAPI (Vector) to stain DNA and reduce fluorescence fading. Images were acquired using an Axio Imager M2 fluorescence microscope (Zeiss, Germany). Image z-stacks were acquired with an Axiocam 512 (Zeiss) monochromatic camera and Apotome 2 (Zeiss); for each image15 z-stack were acquired at 0,5 micrometer Z step. All images were acquired with the same parameters.

Fluorescence signals were quantified using Zen 2.5 Pro software (Zeiss, Germany). Measurements of the eye-antennal disc areas and the ELAV stained areas were performed with the Zen 2.5 Pro software (Zeiss, Germany) on Maximum intensity projections z-stacks. Caspase3 positive foci were also quantified using Zen 2.5 Pro software.

### Evaluation of the eye phenotype

The eye size of RNAi flies was evaluated by comparison with the wild type eye (Oregon-R). Abnormal eyes were assigned to one of the following classes: class 100 that comprises eyes of normal size, and classes 25, 50, and 75 that include eyes with sizes < 25%, < 50%, and < 75% the size of the wild type eye, respectively. Eyes were classified by visual inspection performed independently by at least two researchers. When classification was not clear-cut, the eye was assigned to the higher class in the evaluation of RNAi phenotype without rescue construct (e.g., an eye of dubious class 50 was assigned to class 75), and to the lower class in the presence of a rescue construct.

### Western blot

Brains were dissected in NaCl; protein extracts were separated by SDS-polyacrylamide gel electrophoresis with 10X TRIS-glycine-SDS solution and then transferred on a nitrocellulose membrane. After transfer, membrane was blocked with 3% milk in 25 ml TBS-Tween. Membrane was probed with rabbit anti-GFP (Torrey Pines Biolabs; 1:1000) and mouse anti-tubulin (Sigma-Aldrich;1:7000); secondary antibodies were anti-rabbit 1:5000; anti-mouse 1:5000 conjugated with with the SuperSignal^TM^ West Pico Chemiluminescent Substrate (Thermo Scientific). Images were acquired with Chemidoc (Biorad).

### Transcriptome library preparation and RNA-sequencing

Total RNA was extracted from third instar larval brains or whole third instar larvae with TRIzol reagent (Ambion), treated with AmbionTM DNase I (RNase-free) and extracted with phenol/chloroform using standard protocols. RNA samples were quantified and quality-tested by Tape Station RNA assay. The TruSeq Stranded mRNA kit (Illumina, NGS Padova) was used for library preparation following the manufacturer’s instructions. An rRNA depletion kit was also used. Libraries were then prepared and sequenced on paired-end 150 bp mode on NovaSeq 6000 (Illumina, NGS Padova). Illumina sequencing was performed on both brain and whole larvae RNA samples (in duplicate, listed in Table S2A and S1A) at a depth of 120M. ONT library preparation was performed on RNA extracted from control and mutant brains using the cDNA-PCR SQK-PCB109 kit, to improve novel isoforms detection and reconstruction. This protocol is suitable for full-length transcripts analysis with high throughput, and it is recommended for splice variant and fusion transcripts analysis. RNA from Control and *SmnRNAi* brain samples was ran in parallel using long read ONT sequencing. We performed four runs with MinION, using one flow cell per two samples. Reads coverage and quality information are reported in table S2B.

### Bioinformatic analysis

Raw Illumina data were processed and demultiplexed by Bcl2Fastq 2.20. Adapter sequences were removed with Cutadapt ^101^.

Larval RNA-seq analysis: The Fastq files were used to perform quantification of annotated transcript expression levels with Kallisto v0.44.0^104^ . Differential gene expression analysis was conducted using DEseq2 ^105^ on reference transcriptome (Dmel6.32.106 reference gtf; adjusted p-value<0.05 and ǀ*log2* [Fold Change]ǀ ≥1). Log2[Fold Change] values were shrunk, using apeglm (Generalized linear method), estimator for dispersion^106^ . Gene ontology (GO) analyses on genes with |log2[Fold Change]|≥1 values were performed using DAVID^107^. GO TERMS (BP) with FDR<0.05 were considered significantly enriched (Fisher’s Exact Test for enrichment analysis).Brain RNA-seq analysis:

Raw Illumina data processing and removal of adapter sequences was perfomed as described above. Reads were then mapped to *Drosophila melanogaster* reference genome (Dmel6.32.106) using STAR v2.7.10b ^102^ with parameters *--genomeSAindexNbases 12, --sjdbOverhang 150* for the indexing, and *-- outSAMtype BAM SortedByCoordinate, -- outSAMstrandField intronMotif -- outFilterIntronMotif--RemoveNoncanonical* for the alignment. The resulting BAM files were used to perform the assembly of novel transcripts using Stringtie v2.1.7 ^103^ with parameters *-m 100 -g 50 -c 10* using Dmel6.32.106 reference gtf. We used Kallisto v0.44.0^104^ to quantify annotated transcript expression levels. Differential gene expression analysis and GO analysis were conducted as reported above. Fast5 files produced by ONT sequencing runs were converted to Fastq files using Guppy v6.0.7 basecaller^108^; all the reads with average quality less than 9 were discarded. The Fastq files were aligned to the reference genome using minimap2 v2.17 ^109^ using default parameters, and FLAIR v1.5.1 ^71^ was used to identify novel isoforms (supported by at least 3 ONT reads). Out of all the reads generated from our samples, 70% of them were full-length reads spanning the entire CDS. To generate a single Illumina-ONT extended GTF file containing all novel and annotated isoforms detected by short- and long-read sequencing we used GFFcompare v0.11.2 ^110^. Briefly, the ONT GTF that includes reconstructed ONT isoforms, was merged with the GTF generated using Stringtie and including both annotated and novel isoforms detected by Illumina short-read sequencing. We used *-r -D* options to compare reference annotations and discard duplicates, respectively (i.e., transcripts identified in both ONT and Illumina datasets were counted only once and assigned to the ONT dataset). Transcripts described by this extended GTF were then quantified using Illumina short-reads with Kallisto, considering isoforms with TPM>0.1, generated from genes with TPM>1 in at least one sample. For alternative splicing analysis we used SUPPA2 ^75^ to extract alternative splicing events from the expanded GTF file containing novel and annotated transcripts and to find significantly changed alternative splicing events. For all the seven AS categories (A3, A5, AF, AL, SE, MX, RI), the percentage of splicing inclusion (PSI) was calculated. The significance of differentially spliced events was assessed based on a ǀdPSIǀ≥0.3 and a p-value<0.05 where dPSI represents the difference in PSI between mutant and control samples. For the analysis of transposable elements deregulation we employed the TEtranscripts tool^111^ for the quantification. We mapped short-reads against the reference transcriptome using the STAR parameters --*outFilterMultimapNmax 100 -- outAnchorMultimapNmax 100* and we utilized the TEtranscripts v2.2.3 module for the quantification of annotated TEs using TE annotations file provided by the tool (dm6_BDGP_rmsk_TE.gtf) and reference transcriptome. *Log2*[Fold Change] values calculated by the tool were filtered for ǀlog2[Fold Change]ǀ≥2 and adjusted p-value<0.05.

### RT-qPCR analysis

RNA was extracted by Trizol reagent (Ambion), treated with AmbionTM DNase I (RNase-free), and extracted with phenol/chloroform using standard protocols. The integrity of RNA samples was evaluated by gel electrophoresis. 1 μg of intact RNA (with a 28S:18S rRNA ratio = 2:1) was reverse transcribed using RevertAid H Minus First Strand cDNA Synthesis Kit (Thermo Scientific, EP0451). Real-time PCR reactions were performed using SensiFASTTM SYBR No-ROX Kit, with QuantStudioTM 3 Real-Time PCR System under the following thermal cycling conditions: an initial step of 2 min at 95°C, then 5s denaturation at 95 °C, and 30s elongation at 62 °C. The relative quantification of gene expression was carried out using the 2-ΔΔCt method and the *Rp49* gene for normalization. A total of three experiments were performed for three biological replicates and the significance was assessed by ANOVA. The oligonucleotides used in this study are reported in Table S4.

### HPLC detection and Statistical analysis

Frozen third instar larvae samples (20 to 40 each) were homogenized in 0.2 M TCA (5 μL/larva), sonicated (4 cycles, 15 s each), and centrifuged at 13,000 × g for 20 min. TCA supernatants were then neutralized with NaOH and subjected to pre-column derivatization with o-phthaldialdehyde (OPA)/N-acetyl-L-cysteine (NAC). Diasteroisomer derivatives were resolved on a UHPLC Agilent 1290 Infinity (Agilent Technologies, Santa Clara, CA, USA) using a ZORBAX Eclipse Plus C8, 4.6 × 150 mm, 5 μm (Agilent Technologies, Santa Clara, CA, USA) under isocratic conditions (0.1 M sodium acetate buffer, pH 6.2, 1% tetrahydrofuran, and 1.5 mL/min flow rate). A washing step in 0.1 M sodium acetate buffer, 3% tetrahydrofuran, and 47% acetonitrile was performed after every single run. Identification and quantification of D-Asp, L-Asp, L-Glu, D-Ser, L-Ser, L-Gln, Gly, and GABA were based on retention times and peak areas, compared with those associated with external and internal standards ^39^. All the precipitated protein pellets from larvae samples were solubilized in 1% SDS solution and quantified by bicinchoninic acid (BCA) assay method (Pierce™ BCA Protein Assay Kits, Thermofisher scientific, Rockford, IL, USA). The concentration of amino acids in larvae homogenates was normalized to the total protein content and to the number of larvae per sample and expressed as pmol/mg protein/larva.

Statistical analyses were performed using Prism 8 for Windows version 8.0.2. Amino acid concentrations in the whole larvae were compared using non-parametric Kruskall-Wallis test followed by original false discovery rate (FDR) method of Benjamini and Hochberg’s *post-hoc* for multiple comparisons.

## Data availability

ONT sequencing data: GSE285429 https://www.ncbi.nlm.nih.gov/geo/query/acc.cgi?acc=GSE285429 token for ONT data access: urgtwwgmvnmdlyb

Illumina sequencing data: GSE285430 https://www.ncbi.nlm.nih.gov/geo/query/acc.cgi?acc=GSE285430 token for Illumina data access: ojyfmwumrjoxvon

## Supporting information

Supplementary_material_Scatolini

## Acknowledgements

This work was supported by a grant from AIRC IG 26496 (to G.D.R.)

