## Supplementary_material_Scatolini for "RNase H1 counteracts DNA damage and ameliorates SMN-dependent phenotypes in a *Drosophila* model of Spinal Muscular Atrophy"

**Supplementary Figure S1**

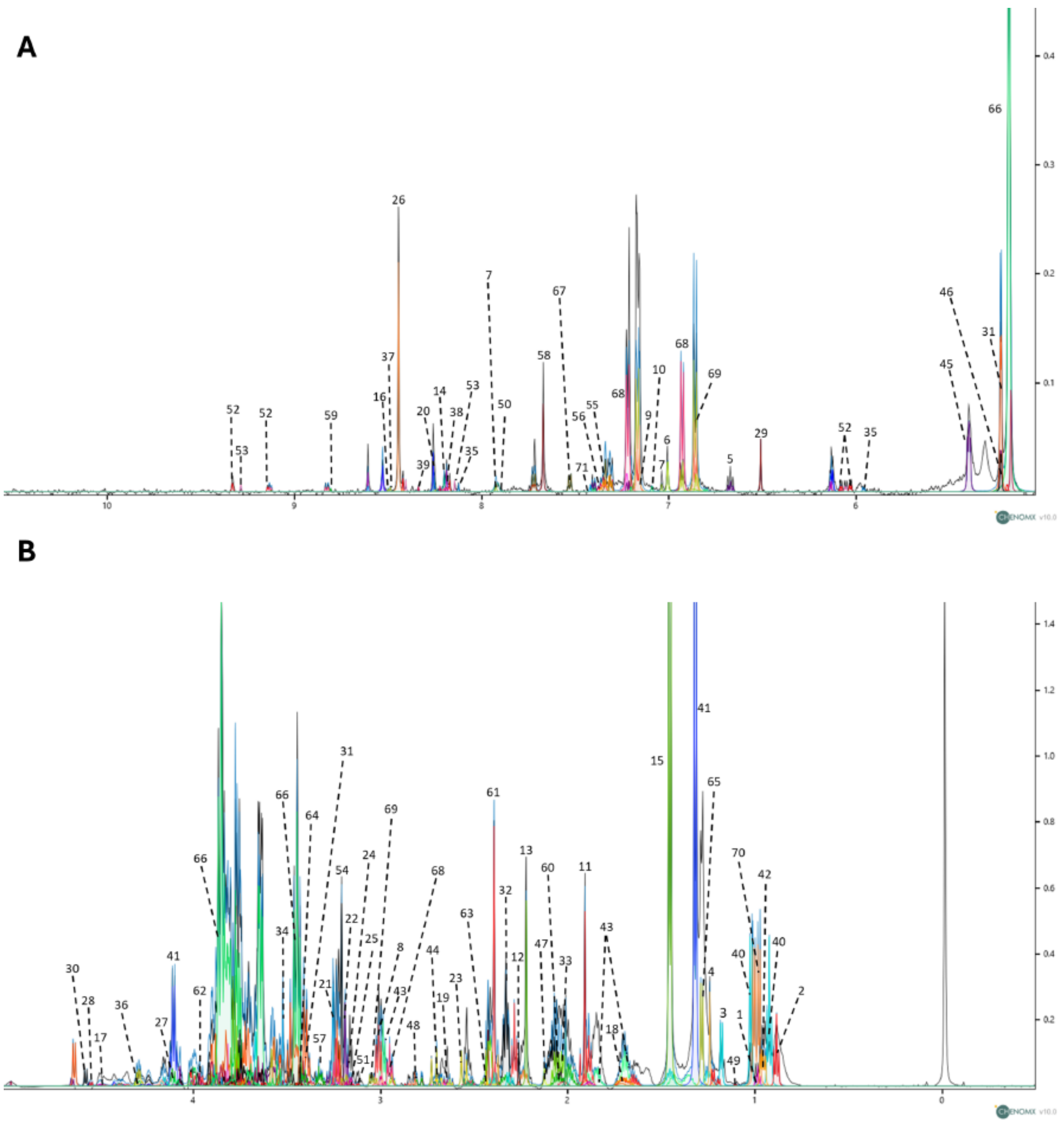


**1**: 2-Aminobutyrate; **2**: 2-Hydroxybutyrate; **3**: 3-Hydroxybutyrate; **4**: 3-Hydroxyisovalerate; **5**: 3-Hydroxykynurenine; **6**: 3-Methoxytyramine; **7**: 3-Methylhistidine; **8**: 4-Aminobutyrate; **9**: 4-Hydroxyphenyllactate; **10**: 5-Hydroxyindole-3-acetate; **11**: Acetate; **12**; Acetoacetate; **13**: Acetone; **14**: Adenine; **15**: Alanine; **16**: AMP; **17**: Anserine; **18**: Arginine; **19**: Aspartate; **20**: ATP; **21**: Betaine; **22**: Choline; **23**: Citrate; **24**: Creatinine; **25**: Cysteine; **26**: Formate; **27**: Fructose; **28**: Fucose; **29**: Fumarate; **30**: Galactose; **31**: Glucose; **32**: Glutamate; **33**: Glutamine; **34**: Glycine; **35**: GTP; **36**: Guanidinosuccinate; **37**: Histydine; **38**: IMP; **39**: Inosine; **40**: Isoleucine; **41**: Lactate; **42**: Leucine; **43**: Lysine; **44**: Malate; **45**: Maltose; **46**: Mannose; **47**: Methionine; **48**: Methylguanidine; **49**: Methylsuccinate; **50**: N-Acetylaspartate; **51**: N-Acetyl-L-tyrosine; **52**: NAD+; **53**: NADP+; **54**: O-Phosphocholine; **55**: Phenylacetate; **56**: Phenylalanine; **57**: Proline; **58**: Pyridoxine; **59**: Pyrimidine; **60**: Pyroglutamate; **61**: Pyruvate; **62**: Serine; **63**: Succinate; **64**: Taurine; **65**: Threonine; **66**: Trehalose; **67**: Tryptophan; **68**: Tyramine; **69**: Tyrosine; **70**: Valine; **71**:1-Methylhistidine.

#### Supplementary Figure S2

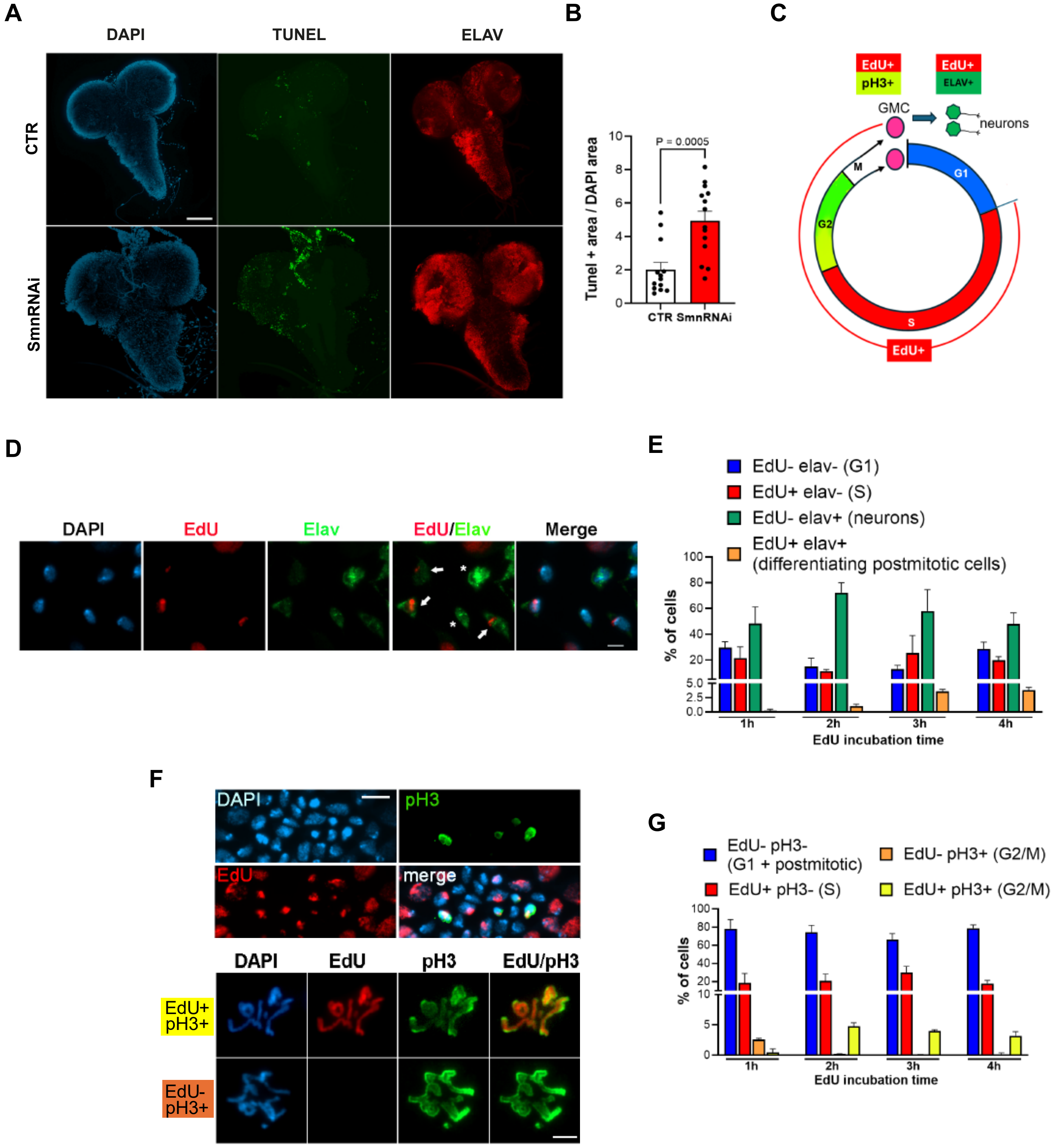

#### Supplementary Figure S3

**A**

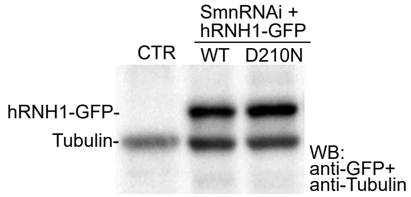


**B**

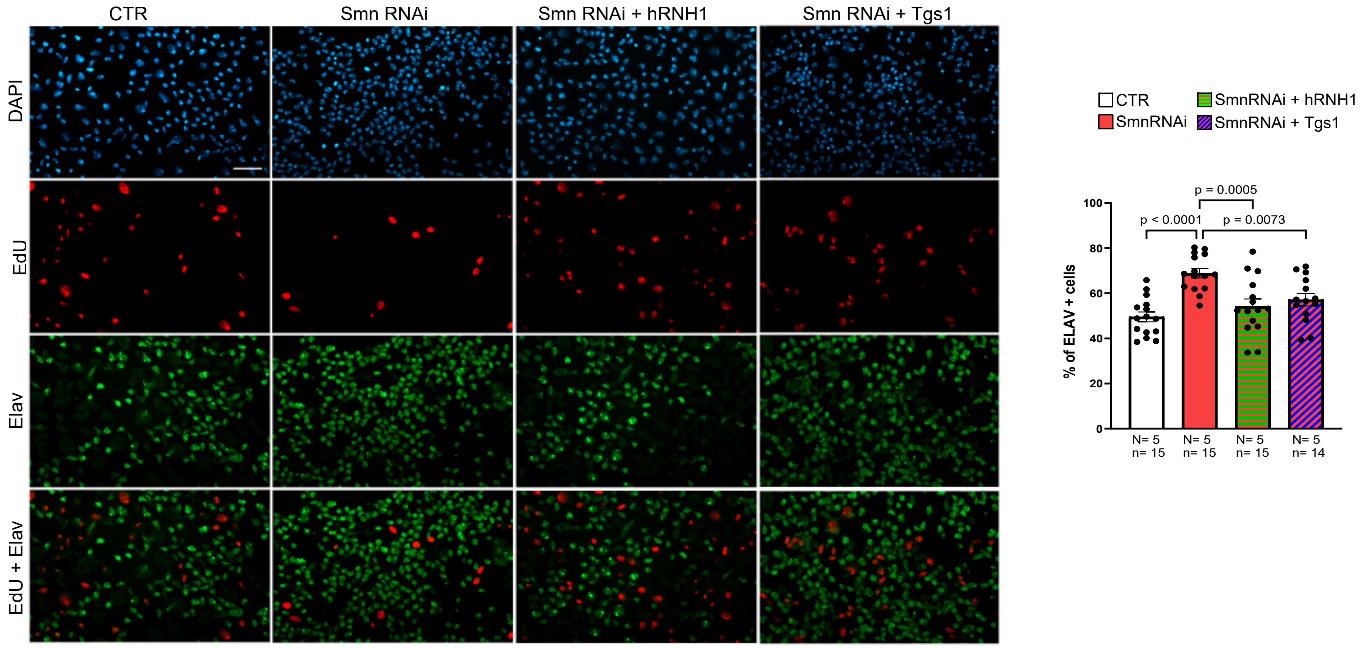


**C**

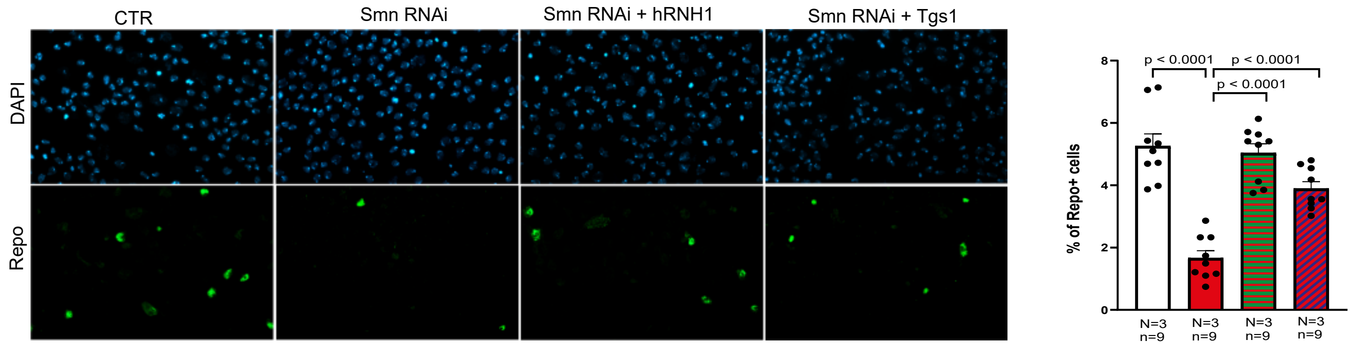

#### Supplementary Figure S4

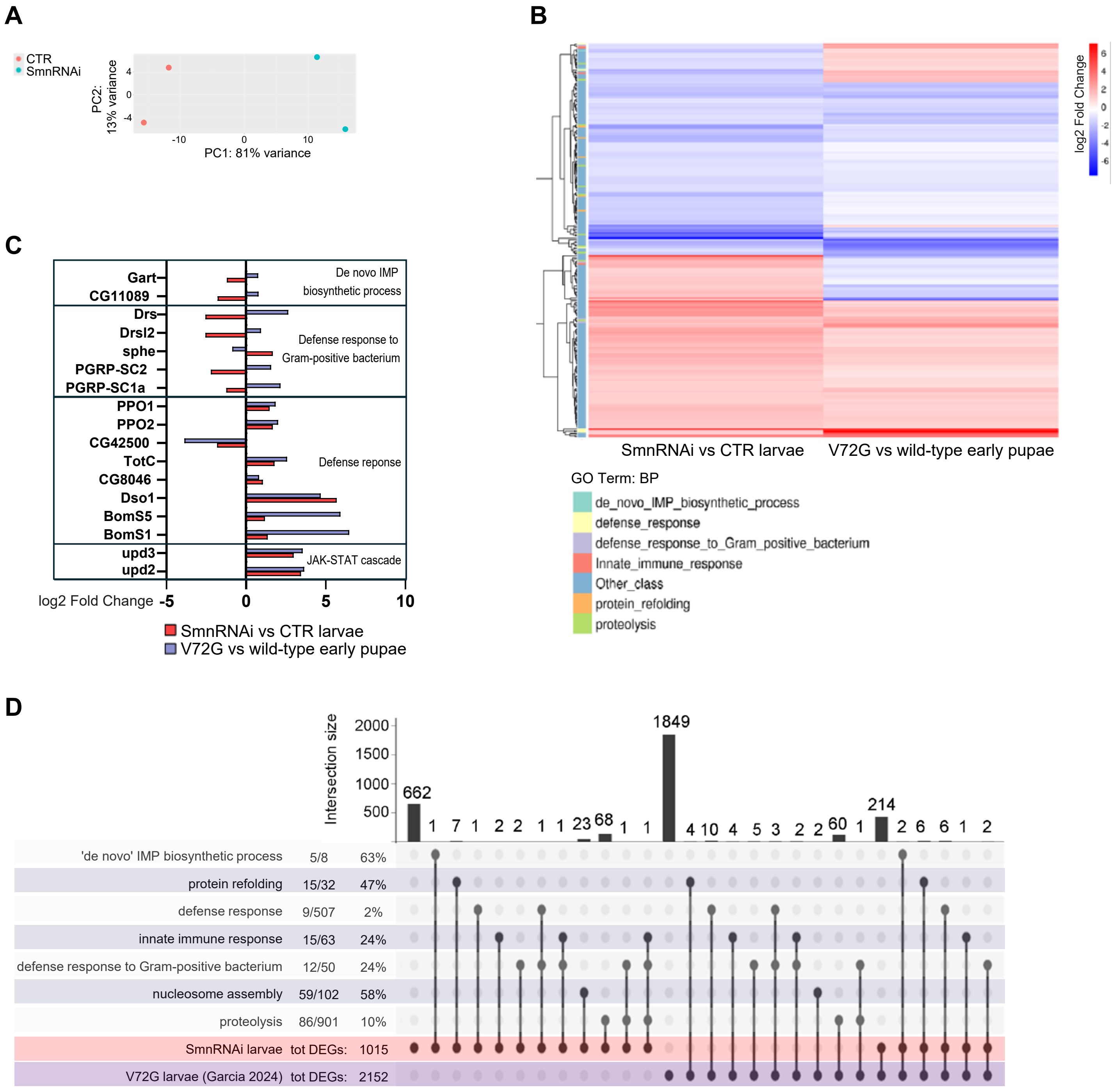

### Supplementary Figure S5

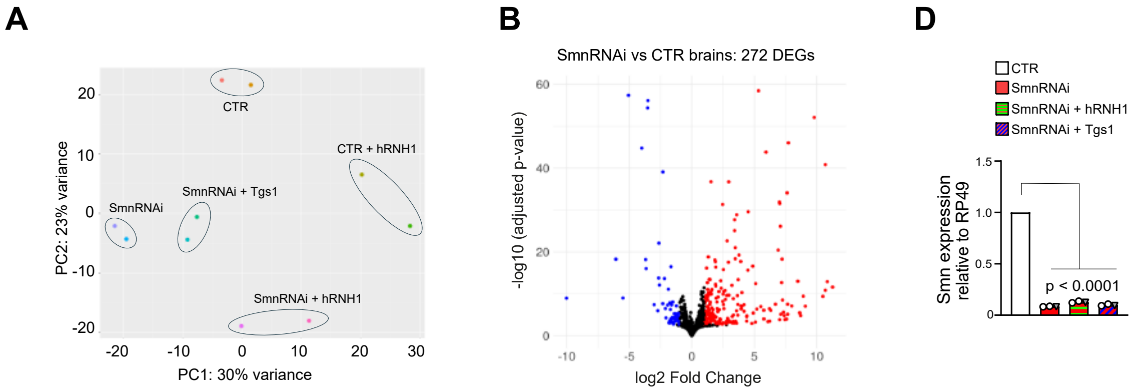


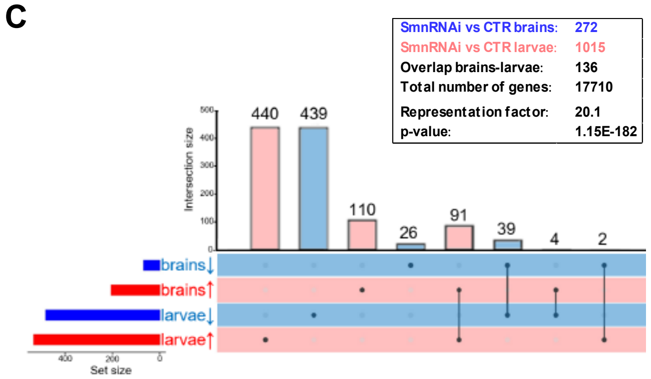


E

| | Retrotransposons transcripts log2FoldChange | |
|---|---|---|
| | LARVAE | BRAINS |
| transposable element | SmnRNAi vs CTR | |
| Copia | 2.17 | 3.94 |
| Blastopia | 2.56 | 2.45 |
| Invader4 | 4.32 | 3.83 |
| Invader5 | 2.40 | 1.91 |
| Gypsy | 3.69 | 1.81 |

### Supplementary Figure S6

- SmnRNAi vs CTR
- SmnRNAi + Tgs1 vs CTR
- SmnRNAi vs SmnRNAi + TGS1
- SmnRNAi + hRNH1 vs CTR hRNH1
- SmnRNAi vs SmnRNAi + hRNH1

| | |
|---|---|
| a-rescued by Tgs1 | 48 |
| b-partially rescued by Tgs1 | 7 |
| c- NON rescued | 62 |
| d-undetermined | 25 |

| | |
|---|---|
| a-rescued by hRNH1 | 22 |
| b-partially rescued by hRNH1 | 3 |
| c- NON rescued | 58 |
| d-undetermined | 59 |

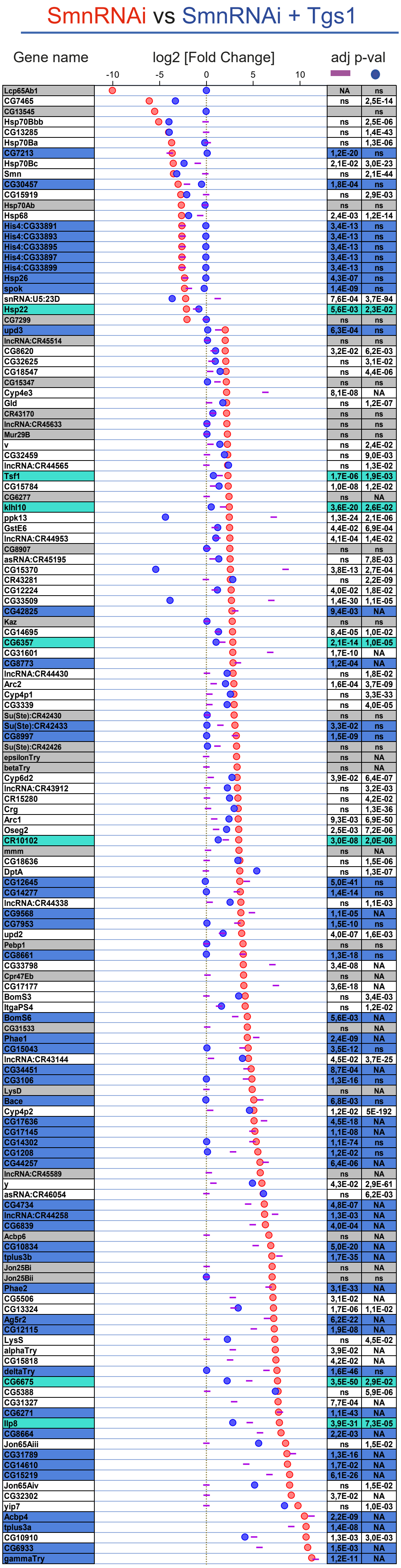


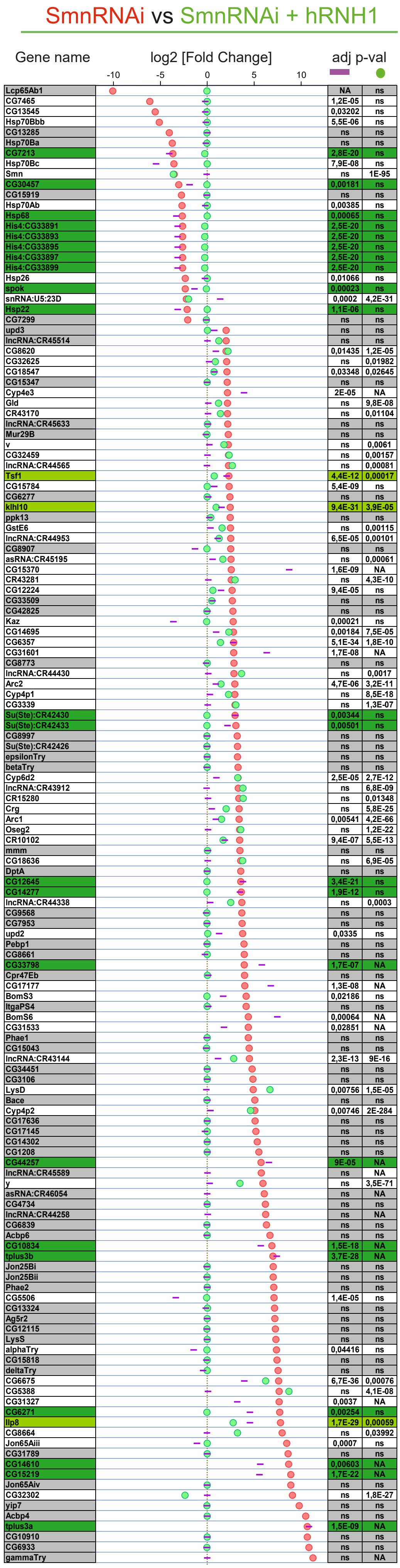

#### Supplementary Figure S7

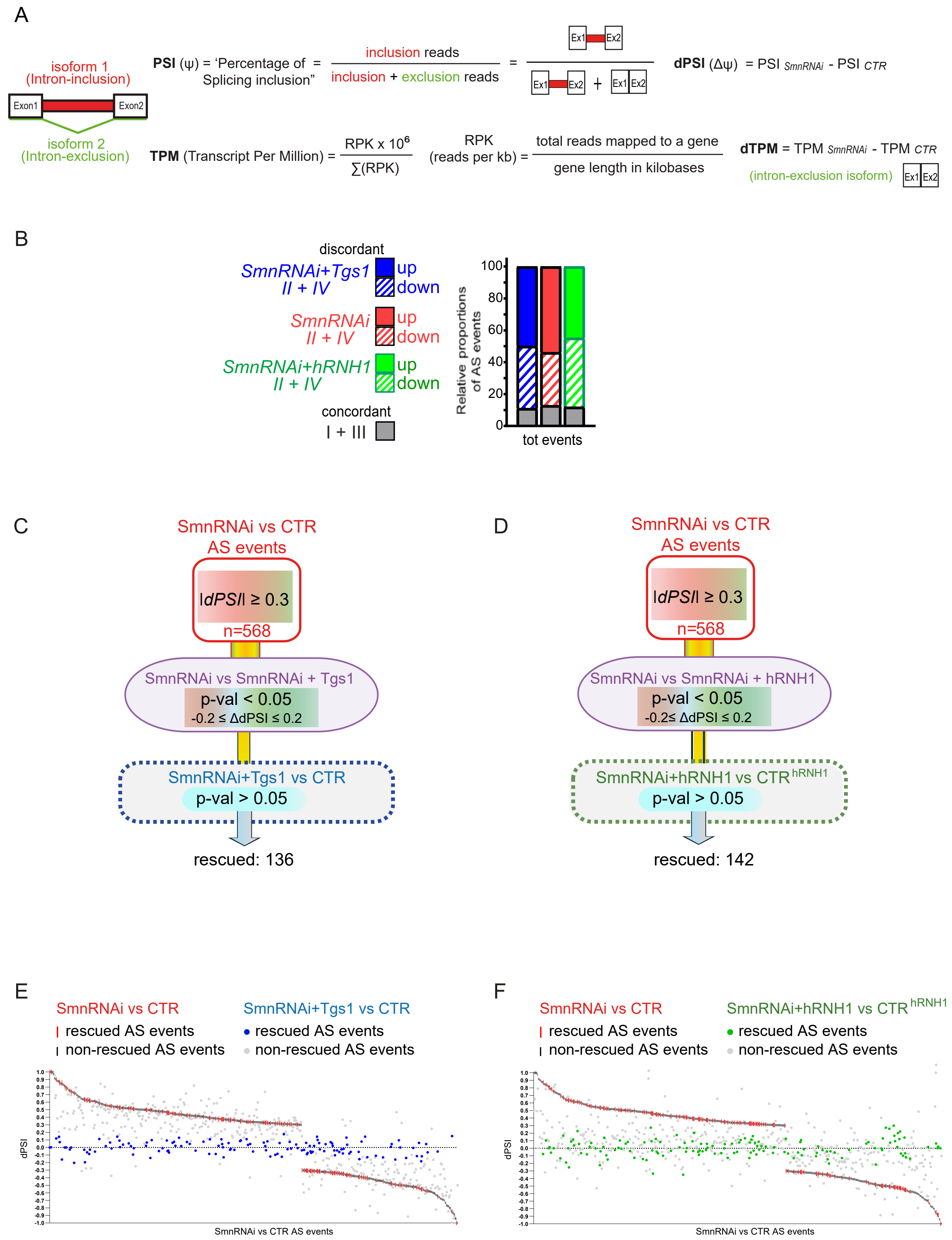

### Supplementary Figure S8

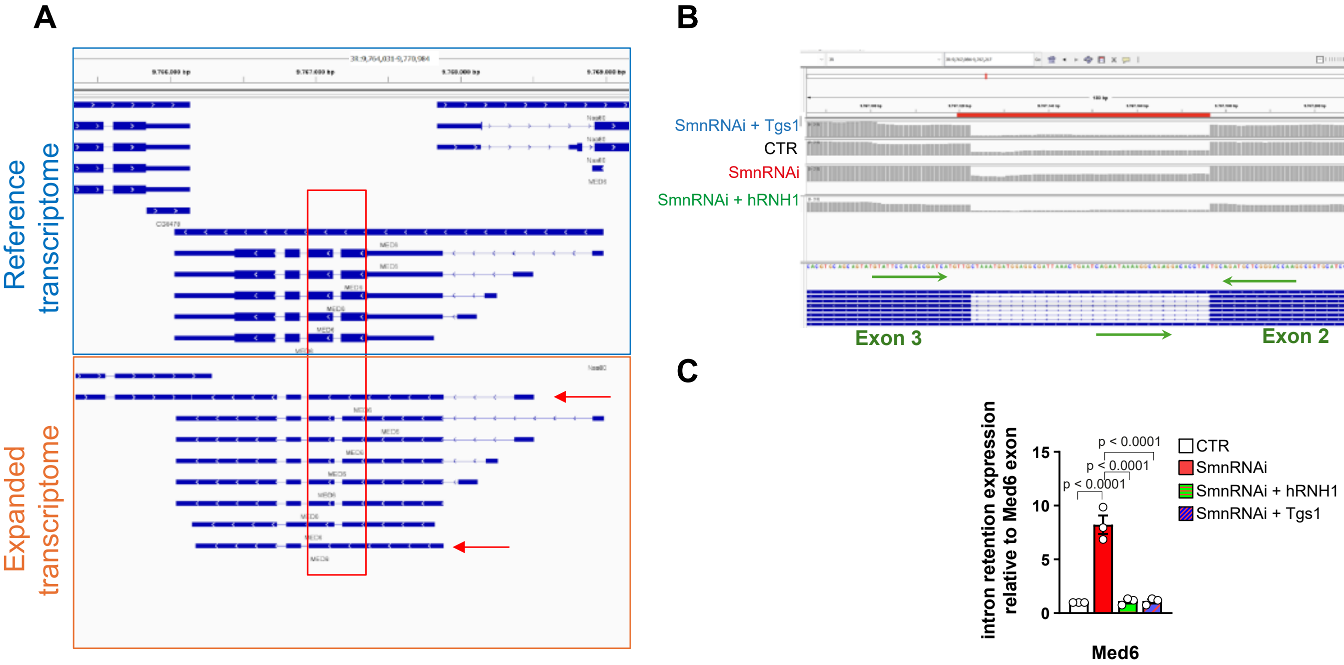

### Supplementary Figure S9

**A**

| ens_gene | ext_gene | log2[Fold Change] | adjusted p-value | KO number | beta-alanine metabolic process | Glutamate GABA | Glutathione | Gly Ser Thr | purine metabolic process | Tyr DOPA | Butanoate metabolism |
|---|---|---|---|---|---|---|---|---|---|---|---|
| FBgn0063494 | GstE6 | 2,34 | 1,13E-12 | K00799 | | | * | | | | |
| FBgn0051343 | CG31343 | 2,26 | 8,94E-08 | K11140 | | | * | | | | |
| FBgn0003961 | Uro | 2,12 | 2,45E-14 | K00365 | | | | | * | | |
| FBgn0038105 | yellow-f2 | 1,94 | 1,81E-04 | K22203 | | | | | | * | |
| FBgn0036992 | Hpd | 1,88 | 1,82E-05 | K00457 | | | | | | * | |
| FBgn0039656 | CG11951 | 1,66 | 3,36E-06 | K11140 | | | * | | | | |
| FBgn0010043 | GstD7 | 1,56 | 0,023 | K00799 | | | * | | | | |
| FBgn0267408 | AOX1 | 1,48 | 2,67E-15 | K00106 | | | | | * | | |
| FBgn0283437 | PPO1 | 1,48 | 6,93E-07 | K00505 | | | | | | * | |
| FBgn0000075 | amd | 1,21 | 0,015 | K01618 | | | | | | * | |
| FBgn0026565 | Ass | 1,18 | 0,004 | K01940 | | * | | | | | |
| FBgn0051233 | CG31233 | 1,13 | 0,001 | K11140 | | | * | | | | |
| FBgn0043025 | Adgf-A2 | 1,07 | 0,045 | K19572 | | | | | * | | |
| FBgn0038020 | GstD9 | 1,04 | 7,58E-06 | K00799 | | | * | | | | |
| FBgn0027493 | AdSS | 1,02 | 1,30E-04 | K01939 | | * | | | * | | |
| FBgn0063495 | GstE5 | 1,02 | 0,011 | K00799 | | | * | | | | |
| FBgn0050446 | Tdc2 | 0,91 | 0,013 | K22329 | | | | | | * | |
| FBgn0264815 | Pde1c | 0,91 | 9,06E-08 | K13755 | | | | | * | | |
| FBgn0010039 | GstD3 | 0,86 | 1,08E-05 | K00799 | | | * | | | | |
| FBgn0030484 | GstT4 | 0,76 | 0,001 | K00799 | | | * | | | | |
| FBgn0003189 | r | 0,67 | 3,18E-05 | K11540 | | * | | | | | |
| FBgn0025837 | CG17636 | 0,58 | 0,033 | K18592 | | | * | | | | |
| FBgn0024150 | Ac78C | 0,56 | 0,002 | K08048 | | | | | * | | |
| FBgn0046114 | Gclm | 0,53 | 0,009 | K11205 | | | * | | | | |
| FBgn0011703 | RnrL | 0,48 | 0,001 | K10807 | | | * | | * | | |
| FBgn0030796 | CG4829 | 0,47 | 0,001 | K18592 | | | * | | | | |
| FBgn0261625 | GLS | 0,46 | 0,029 | K01425 | | * | | | | | |
| FBgn0032775 | CG17544 | 0,39 | 0,040 | K00232 | * | | | | | | |
| FBgn0037684 | Srr | 0,37 | 0,006 | K01754 | | | | * | | | |
| FBgn0038349 | AOX3 | 0,36 | 0,012 | K00106 | | | | | * | | |
| FBgn0000479 | dnc | 0,27 | 0,027 | K13293 | | | | | * | | |
| FBgn0005626 | ple | -0,20 | 0,028 | K00501 | | | | | | * | |
| FBgn0028479 | Mtpalpha | -0,23 | 0,026 | K07515 | * | | | | | | * |
| FBgn0000454 | Dip-B | -0,26 | 0,030 | K01255 | | | * | | | | |
| FBgn0038742 | Arc42 | -0,32 | 0,031 | K00248 | * | | | | | | * |
| FBgn0036099 | CG11811 | -0,32 | 0,027 | K00942 | | | | | * | | |
| FBgn0010548 | Aldh-III | -0,34 | 0,001 | K00129 | * | | | | | * | |
| FBgn0001149 | GstD1 | -0,35 | 0,001 | K00799 | | | * | | | | |
| FBgn0036857 | Aldh7A1 | -0,42 | 0,012 | K14085 | * | | | * | | | |
| FBgn0003308 | ry | -0,44 | 0,016 | K00106 | | | | | * | | |
| FBgn0013307 | Odc1 | -0,49 | 0,001 | K01581 | | | * | | | | |
| FBgn0052626 | AMPdeam | -0,53 | 2,94E-07 | K01490 | | | | | * | | |
| FBgn0031117 | GstT3 | -0,54 | 0,001 | K00799 | | | * | | | | |
| FBgn0261446 | CG13377 | -0,57 | 0,000519224 | K00019 | | | | | | | * |
| FBgn0001125 | Got2 | -0,58 | 0,001 | K14455 | | * | | | | * | |
| FBgn0000150 | awd | -0,59 | 1,27E-06 | K00940 | | | | | * | | |
| FBgn0032729 | L2HGDH | -0,60 | 0,006645548 | K00109 | | | | | | | * |
| FBgn0000109 | Aprt | -0,61 | 0,012 | K00759 | | | | | * | | |
| FBgn0034354 | GstE11 | -0,63 | 0,002 | K00799 | | | * | | | | |
| FBgn0039109 | CG10365 | -0,63 | 0,018 | K07232 | | | * | | | | |
| FBgn0000566 | Cth | -0,65 | 4,75E-05 | K01758 | | | | * | | | |
| FBgn0033381 | GstE13 | -0,65 | 0,005 | K00799 | | | * | | | | |
| FBgn0000052 | Pfas | -0,66 | 1,33E-05 | K01952 | | | | | * | | |
| FBgn0004057 | Zw | -0,66 | 0,021 | K00036 | | | * | | | | |
| FBgn0020389 | Papss | -0,67 | 1,53E-07 | K13811 | | | | | * | | |
| FBgn0051183 | CG31183 | -0,67 | 3,68E-05 | K12323 | | | | | * | | |
| FBgn0038172 | Adgf-D | -0,75 | 7,82E-08 | K19572 | | | | | * | | |
| FBgn0023023 | CRMP | -0,80 | 0,001 | K01464 | * | | | | | | |
| FBgn0036787 | CG4306 | -0,81 | 2,49E-06 | K00682 | | | * | | | | |
| FBgn0035438 | PHGPx | -0,82 | 5,80E-13 | K00432 | | | * | | | | |
| FBgn0034225 | veil | -0,82 | 3,99E-06 | K19970 | | | | | * | | |
| FBgn0032287 | CG6415 | -0,84 | 0,002 | K00605 | | | | * | | | |
| FBgn0004654 | Pgd | -0,85 | 3,08E-11 | K00033 | | | * | | | | |
| FBgn0037999 | CG4860 | -0,88 | 0,004 | K00248 | * | | | | | | * |
| FBgn0036337 | Adk2 | -0,88 | 6,23E-19 | K00856 | | | | | * | | |
| FBgn0270926 | AsnS | -0,88 | 2,32E-05 | K01953 | | * | | | | | |
| FBgn0037513 | pyd3 | -0,90 | 1,75E-05 | K01431 | * | | | | | | |
| FBgn0036030 | Prps | -0,92 | 2,22E-23 | K00948 | | | | | * | | |
| FBgn0063493 | GstE7 | -0,92 | 2,86E-04 | K00799 | | | * | | | | |
| FBgn0003204 | ras | -0,93 | 8,98E-06 | K00088 | | | | | * | | |
| FBgn0052549 | Nt5b | -0,99 | 5,97E-12 | K01081 | | | | | * | | |
| FBgn0041194 | Prat2 | -1,01 | 8,35E-17 | K00764 | | * | | | * | | |
| FBgn0035904 | GstO3 | -1,01 | 3,85E-10 | K00310 | | | * | | | | |
| FBgn0000055 | Adh | -1,09 | 3,54E-05 | K00129 | | | | | | * | |
| FBgn0027945 | ppl | -1,10 | 1,22E-09 | K01079 | | | | * | | | |
| FBgn0014869 | Pglym78 | -1,11 | 8,13E-07 | K01834 | | | | * | | | |
| FBgn0029924 | CG4586 | -1,13 | 5,63E-13 | K00232 | * | | | | | | |
| FBgn0051075 | CG31075 | -1,20 | 4,97E-27 | K00128 | * | | | | | | |
| FBgn0038467 | AdSL | -1,21 | 1,51E-17 | K01756 | | * | | | * | | |
| FBgn0000053 | Gart | -1,21 | 4,42E-06 | K11787 | | | | | * | | |
| FBgn0036975 | CG5618 | -1,22 | 1,08E-04 | K18966 | * | | | | | | |
| FBgn0050104 | NT5E-2 | -1,31 | 1,40E-26 | K19970 | | | | | * | | |
| FBgn0020513 | Paics | -1,33 | 8,07E-75 | K01587 | | | | | * | | |
| FBgn0037370 | CG1236 | -1,35 | 1,98E-28 | K00049 | | | | * | | | |
| FBgn0035348 | CG16758 | -1,57 | 9,86E-17 | K03783 | | | | | * | | |
| FBgn0001145 | Gs2 | -1,61 | 9,18E-08 | K01915 | | * | | | | | |
| FBgn0014427 | CG11899 | -1,61 | 1,16E-05 | K00831 | | | | * | | | |
| FBgn0063497 | GstE3 | -1,64 | 3,19E-27 | K00799 | | | * | | | | |
| FBgn0001142 | Gs1 | -1,71 | 1,63E-23 | K01915 | | * | | | | | |
| FBgn0032350 | CG6287 | -1,78 | 1,05E-11 | K00058 | | | | * | | | |
| FBgn0039241 | CG11089 | -1,78 | 5,79E-13 | K00602 | | | | | * | | |
| FBgn0041710 | yellow-f | -1,92 | 9,12E-18 | K22203 | | | | | | * | |
| FBgn0029823 | Shmt | -1,96 | 1,34E-74 | K00600 | | | | * | | | |
| FBgn0037801 | CG3999 | -2,06 | 1,88E-27 | K00281 | | | | * | | | |
| FBgn0000153 | b | -2,17 | 9,56E-05 | K18966 | * | | | | | | |
| FBgn0022774 | Oat | -2,65 | 9,36611E-28 | K00819 | | * | | | | | |
| FBgn0263236 | SP1029 | -2,88 | 1,21E-06 | K11140 | | | * | | | | |
| FBgn0000422 | Ddc | -3,36 | 1,25E-04 | K01593 | | | | | | * | |
| FBgn0046253 | CG3502 | -4,24 | 1,58E-12 | K11140 | | | * | | | | |

**B**

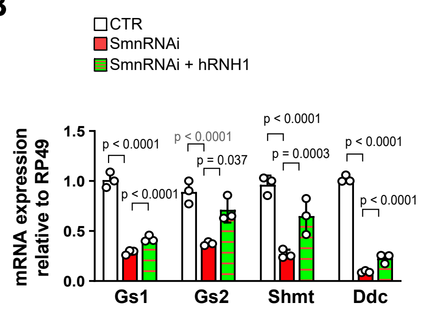

**Supplementary Figure legends**

**Figure S1.** Representative 1D $^1$H NOESY spectrum of extracts from *SmnRNAi* and CTR third instar larvae**.**

**Figure S2.** ***SmnRNAi*** **brains accumulate DNA damage**
(A) *SmnRNAi* larval brains were stained with TUNEL (green) to mark cells with fragmented DNA, and for ELAV (red) that labels differentiated cells. Compared to control brains (CTR), the number of cells marked by TUNEL is clearly increased. Nuclei were counterstained with DAPI (blue); Size bar: 100μM
(B) Quantification of TUNEL signals. p values: two-tailed unpaired t-test, df=25. 13 brains examined per genotype.

(C) Schematic of the cell cycle depicting the last cell division that generates two ganglion mother cells showing EdU incorporation and pH3 expression prior to expression of ELAV in postmitotic neurons.

(D-G) Immunofluorescence analysis performed on larval brains incubated with EdU for the indicated time intervals. Following the click-it reaction, for detection of EdU labeling, brains were immunostained for the neuronal marker ELAV (D-E), or the mitotic marker pH3 (F-G). 2-hours EdU pulsing time resulted in a nearly complete labeling of the population of cells that had entered S phase during the incubation time and had proceeded through mitosis, i.e., 97% pH3 positive cells had also incorporated EdU (quantified in G). In these conditions, only a few ELAV+ postmitotic cells, were also EdU+. Longer incubation times (≥ 3 hours), show a gradual increase in the proportion of postmitotic neurons labeled with EdU (ELAV+EdU+, quantified in E). This suggests that within the initial 2-hour pulse, EdU+ ELAV- cells represented the cycling cell population (S/G2/M phase), while the EdU- ELAV- population mainly consists of cells in the G1 phase or in a quiescent state. This EdU- ELAV- population might also include a small proportion of cells that have completed DNA synthesis and/or mitosis just before the EdU pulse and that are not matured into neurons yet. >1300 wild type cells examined per condition.

**Figure S3. Alterations of the cell cycle profiles induced by Smn depletion are rescued by hRNH1 or Tgs1**
(A) Western blot, probed for GFP, on brains of larvae constitutively expressing the hRNH1-GFP or catalytically dead hRNH1$^{D210N}$-GFP, (under control of the Tubulin promoter),

showing that hRNH1WT-GFP and hRNH1 hRNH1$^{D210N}$ -GFP are expressed at comparable levels. Tubulin was used as loading control.
(B) Brains with the indicated genotypes were incubated with EdU for 2h, immunostained for ELAV, and counterstained with DAPI. Size bar in (B, C): 20μM
(C) Brains with the indicated genotypes were immunostained for Repo and counterstained with DAPI.
The histogram bars on the right panels represent the quantifications of the cells labeled by ELAV (in B) and REPO (in C). IF-positive cells were automatically quantified with the Zeiss Zen software. p values: one-way ANOVA, $F(3, 55) = 10.11$, $p<0.0001$ (B), $F(3, 32) = 31.42$, $p<0,0001$ (C). N: number of experiments; n: number of brains; cells examined per sample: > 26,000 (B); >14,000 (C)

**Figure S4. Differentially expressed genes in *SmnRNAi* larvae**

(A) Principal Component Analysis (PCA) plot showing the clustering of *SmnRNAi* and control larvae samples based on their gene expression profile measured via short-read RNA-Seq. Principal component 1 (PC1) and 2 (PC2) were identified based on variance-stabilizing transformed (VST) counts relative to the 1000 most variable genes.
(B) Heatmap illustrating the comparison between the log2[Fold Change] values for common differentially expressed genes in *SmnRNAi* mutant larvae (this paper) and *Smn*$^{V72G}$ early pupae reported in [30, 66], annotated by Gene Ontology (GO) biological process term (FDR <0.05).
(C) Comparison of the expression levels of genes differentially expressed in both *SmnRNAi* and *Smn*$^{V72G}$ mutant RNA datasets that fall within the GO terms of interest, including de novo IMP biosynthetic process, defense response to Gram positive bacterium and defense response.
(D) UpSet plot illustrating the overlap of differentially expressed (DE) genes between *SmnRNAi* and *Smn*$^{V72G}$ datasets, with their distribution across significant GO biological process categories. Values on the left indicate gene fractions and their corresponding percentages relative to the size of each GO category. The gene overlap between the two datasets was evaluated using a hypergeometric test ($p<1.359e-56$, representation factor = 2.9).

**Figure S5. Differential gene expression analysis in *SmnRNAi* brains and larvae**

(A) Principal Component Analysis (PCA) plot showing the clustering of the indicated *SmnRNAi* and control brain samples based on their gene expression profile measured via RNA-Seq. Principal component 1 (PC1) and 2 (PC2) were identified based on variance-stabilizing transformed (VST) counts relative to the 1000 most variable genes.
(B) Volcano plot of DEGs in *SmnRNAi* vs control brains (Blue and red dots: |log2[Fold Change]|≥1 and adjusted p-value<0.05)
(C) UpSet plot representing the comparison between significant up- (red) or down- (blue) regulated genes (|log2[Fold Change]|≥1 and adjusted p-value<0.05) identified in *SmnRNAi* brains and whole larvae. The overlap between the two RNA datasets was calculated by hypergeometric test (p<1.153e-182, representation factor 20.1)
(D) RT-qPCR analysis of RNA from larvae with the indicated genotypes, demonstrating comparable knockdown efficiency of the *Smn* transcript in *SmnRNAi* samples relative to the control. Bars represent the fold change of *Smn* transcript levels normalized to *Rp49*. Data are relative to control flies (set to 1). p-values one-way ANOVA, F (3, 8) = 13719; p<0.0001.
(E) Table showing the log2[Fold Change] values of the indicated retrotransposon (TE) transcripts in *SmnRNAi* brains and larvae datasets compared to controls (see also Table S2H,S1E).

**Figure S6. Differentially expressed genes in *SmnRNAi, SmnRNAi* + *Tgs1* and *SmnRNAi* + *hRNH1 larval brains***

Graphical representation of the log2[Fold Change] values for 142 greatly varying DEGs in *SmnRNAi vs. control* brains. Color-coded symbols indicate the log2[Fold Change] of DEGs in pairwise comparisons between the indicated conditions: (i) *SmnRNAi vs. control* (red dots); (ii) *SmnRNAi* + *rescue construct vs. control* (Tgs1 - blue dots or hRNH1 - green dots); (iii) *SmnRNAi vs. SmnRNAi* + *rescue construct* (purple dashes).

DEGs were categorized into four classes (a to d) based on their expression status in each condition. A gene identified as differentially expressed in "*SmnRNAi vs. control*" brains is considered 'rescued' upon the expression of either Tgs1 or hRNH1 (and assigned to class a) if the following three criteria are met (see also Table S2G):

1. The gene must be differentially expressed in the "*SmnRNAi vs. control*" comparison (represented by red dots; |log2[Fold Change]| ≥ 2 and adjusted p-value<0.05);

2. The same gene must be non-differentially expressed (adjusted p-value>0.05) in the "*SmnRNAi + rescue construct vs. control*" comparison (blue dots for *SmnRNAi* + Tgs1, green dots for *SmnRNAi* + *hRNH1*);
3. The log2[Fold Change] for the gene must be statistically significant in the comparison of "*SmnRNAi vs. SmnRNAi + rescue construct*", and its absolute log2[Fold Change] value should be close to that observed in the "*SmnRNAi vs. control*" comparison (represented by purple dashes).Treshold values are reported in Table S2G.

Genes with significant log2[Fold Change] values (adjusted p-value<0.05) in the "*SmnRNAi + rescue construct vs. control" comparison*, but with lower absolute values than the log2[Fold Change] in the "*SmnRNAi vs. control*" comparison, were considered partially rescued and assigned to class b.

**Figure S7. Characterization of AS events altered in SmnRNAi, *SmnRNAi* + *Tgs1* and *SmnRNAi* + *hRNH1* brains**

(A) Definition of the parameters described in the scatter plot in Figure 7B.

(B) Distribution of differential alternative splicing events partitioned into II and IV quadrants across the seven major classes (A3, A5, AF, AL, MX, RI, SE) in *SmnRNAi* (red bars), *SmnRNAi* + *Tgs1* (blue bars), and *SmnRNAi* + *hRNH1* (green bars) larval brains. The grey bars indicate the proportions of events falling within the I and III quadrants (dark grey) across the three comparisons, reflecting concordant changes in the dPSI and dTPM values of the isoform with and without the AS inclusion event. These concordant events were excluded from our analyses to rule out altered AS events that were merely a consequence of varying transcriptional activity at that locus. Thus, we focused exclusively on events where the expression level changes of the isoforms with and without the AS inclusion event were discordant; specifically, events demonstrating upregulation of the isoform with the inclusion alongside downregulation of the isoform with exclusion (IV quadrant in Figure 7B) and vice versa (II quadrant). The proportion of discordant AS events (IV or II quadrant) was about 80% in all three datasets.
(C-D) Workflow outlining the assessment of 568 alternative splicing (AS) events significantly altered in *SmnRNAi brains* (│dPSI│≥0.3; adjusted p-value<0.05) and rescued

in *SmnRNAi* + *Tgs1* (C) or *SmnRNAi* + *hRNH1* (D) samples. Each AS event was evaluated through pairwise comparisons based on the following criteria:

i) the rescued AS event must show significant alteration in the *“mutant vs. mutant* + *rescue construct”* comparison (adjusted p-value<0.05)

ii) the difference in dPSI values (ΔdPSI) must be smaller than 0.2 (−0.2 ≤ ΔdPSI ≤ 0.2), where

$$\Delta dPSI = dPSI^{\,SmnRNAi \text{ vs CTR}} - dPSI^{\,SmnRNAi \text{ vs } SmnRNAi \text{ + rescue construct}}$$

iii) The rescued event should not show significant alteration in dPSI in the *“mutant + rescue construct vs CTR”* comparison (adjusted p-value >0.05). Events with $\left| dPSI^{\,SmnRNAi \text{ + rescue construct vs CTR}} \right| < 0.2$ and adjusted p-value>0.05 were also considered rescued. See also Table S3B.

(E) The graph displays dPSI values for 568 significant alternative splicing (AS) events in the “*SmnRNAi vs. CTR”* comparison (red and black dashes) paired with dPSI values in the “*SmnRNAi* + *Tgs1 vs. CTR”* comparison (blue/grey dots). Significance was assessed using the two-tailed Wilcoxon matched-pairs signed rank test (p<0.0001) and two-tailed Spearman’s correlation (ρ = 0.7223). Red and blue symbols indicate 136 events with significantly rescued dPSI values in the “*SmnRNAi* + *Tgs1 vs CTR”* comparison (blue dots) compared to “*SmnRNAi vs CTR*” (red dashes). Black and grey symbols represent events with no significant difference between the two comparisons: “*SmnRNAi* + *Tgs1 vs CTR”* (grey circles) and “*SmnRNAi vs CTR*” (black dashes), indicating no rescue by Tgs1 expression. AS events were filtered based on dPSI≥0.3. For more details on rescue criteria, see panel C and Table S3B.

(F) The graph displays dPSI values for 568 significant alternative splicing (AS) events in the “*SmnRNAi vs. CTR”* comparison (red and black dashes) paired with dPSI values in the “*SmnRNAi* + *hRNH1 vs. CTR”* comparison (green/grey dots). Significance was assessed using the two-tailed Wilcoxon matched-pairs signed rank test (p<0.0001) and two-tailed Spearman’s correlation (ρ = 0.3790). Red and green symbols indicate 142 events with significantly rescued dPSI values in the “*SmnRNAi* + *hRNH1 vs. CTR”* comparison (green circles) compared to “*SmnRNAi vs CTR*” (red dashes). Black and grey symbols represent events with no significant difference between the two comparisons: “*SmnRNAi* + *hRNH1 vs CTR”* (grey dots) and “*SmnRNAi vs CTR*” (black dashes), indicating no rescue by hRNH1 expression. AS events were filtered based on dPSI≥0.3. For more details on rescue criteria, see panel D and Table S3B.

**Figure S8. Characterization of intron retention events in *SmnRNAi*, *SmnRNAi* + *Tgs1* and *SmnRNAi* + *hRNH1* brains**

(A) Visualization of the intron retention event in the *Med6* gene using IGV (Integrative Genome Viewer). The intron retention event, indicated by the arrows, is present in the Illumina-ONT expanded transcriptome (bottom), but not in the reference transcriptome (top).
(B) Illumina short-read between exon 2 and 3 of the *Med6* gene, across different samples, with retained intron highlighted in red. Green arrows indicate the primers used for amplification.
(C) RT-qPCR comparative quantification of intron retention containing isoforms of the *Med6* gene, normalized to the adjacent downstream exons. Data from 3 biological replicates (average of 3 technical replicates) are relative to *Rp49* and are normalized to control brains; p-values: one way-ANOVA, $F(3, 8) = 62.06$; $p<0.0001$.

**Figure S9. List of genes differentially expressed in *SmnRNAi* vs control larvae, which are implicated in SMA-relevant metabolic pathways**
(A) List of genes differentially expressed in *SmnRNAi* compared to control larvae, encoding enzymes involved in amino acid metabolism.
(B) RT-qPCR analysis performed on RNA from larvae with the indicated genotypes showing the differential expression levels of the indicated genes compared to control. Bars represent transcript expression levels (mean with SD) from three biological replicates (average of three technical replicates). Data are normalized to *Rp49* and are relative to control larvae (set to 1). p-values: two-way ANOVA with Tukey's Multiple Comparisons test, genotype effect: $F(2, 24)= 226.4$; $p<0.0001$.

**Supplementary tables description**
**Table S1**. Illumina RNA-seq analysis on control and *SmnRNAi* whole third instar larvae.

**Table S2**. Illumina and ONT RNA-seq analysis on control*, SmnRNAi, SmnRNAi+Tgs1, SmnRNAi+hRNH1* brains.

**Table S3.** Analysis of differential alternative splicing events in brain RNA samples : *CTR, SmnRNAi, SmnRNAi+Tgs1, SmnRNAi+hRNH1*

**Table S4.** List of the oligonucleotides used in this work.
